# Large effect loci have a prominent role in Darwin’s finch evolution

**DOI:** 10.1101/2022.10.29.514326

**Authors:** Erik D. Enbody, Ashley T. Sendell-Price, C. Grace Sprehn, Carl-Johan Rubin, Peter M. Visscher, B. Rosemary Grant, Peter R. Grant, Leif Andersson

**Affiliations:** Department of Medical Biochemistry and Microbiology, Uppsala University, Uppsala, Sweden; Institute for Molecular Bioscience, The University of Queensland, Brisbane, Queensland, Australia; Department of Ecology and Evolutionary Biology, Princeton University, Princeton, NJ, USA; Department of Veterinary Integrative Biosciences, Texas A&M University, College Station, USA

## Abstract

A fundamental goal in evolutionary biology is to understand the genetic architecture of adaptive traits and its evolutionary relevance. Using whole-genome data of 3,958 Darwin’s finches on the Galápagos Island of Daphne Major we identify six loci of large effect that explain 46% of the variation in beak size of *Geospiza fortis*, a key ecological trait. Allele frequency changes across 30 years at these loci affected beak morphology in two ways. An abrupt change in beak size occurred in *Geospiza fortis* as a result of natural selection associated with a drought, and a more gradual change occurred in *G. scandens* as a result of introgressive hybridization. This study demonstrates how large effect loci are a major contributor to the genetic architecture of rapid diversification during adaptive radiations.

**One Sentence Summary:** Allele frequency change at six loci of large effect causes evolutionary change in key ecological traits.

## Main Text

Many traits, like size, shape and its components, vary continuously and are referred to as quantitative traits controlled by quantitative trait loci (QTLs) (Falconer and Mackay 1995, Lynch and Walsh). It is generally assumed that quantitative traits affecting fitness in wild populations have a highly polygenic background (Rockman 2012; Bosse et al. 2017; Barton 2022). If fitness traits are controlled by a huge number of loci each with an infinitesimal small effect, as is the case with most human traits (Yang et al. 2011), then it becomes exceedingly difficult to link selection on phenotypic traits to the adaptive changes at the molecular level (Barton 2022). However, genomic comparisons among species or populations often reveal one or a few isolated loci of large phenotypic effect (Hoekstra et al. 2006; Lamichhaney et al. 2016; Toews et al. 2016; vonHoldt et al. 2018; Hill et al. 2019; Schluter et al. 2021; Semenov et al. 2021), and these have been used to interpret genetic processes during speciation (Seehausen et al. 2014). Yet surprisingly little is known of the effect sizes of loci affecting fitness among individuals within populations (Seehausen et al. 2014), in part due to the large sample sizes of genomic and ecological data needed to quantify them. Effect sizes are needed to improve our understanding of how differences among individuals of a population transform into differences between species. Thus, it is unclear if large effect loci play a prominent role in evolutionary change at least under certain circumstances (Orr and Coyne 1992).

Recent adaptive radiations are particularly suitable for studying the genetic architecture of adaptive traits because the ongoing process of speciation is observable and a link between ecologically relevant traits and population divergence is achievable. For these reasons, adaptive radiations have become a model for identifying genomic loci responsible for ecological divergence among species (Lamichhaney et al. 2015; Campagna et al. 2017; Meier et al. 2018; Marques et al. 2019; McGee et al. 2020). Few wild populations are better suited to investigate the link between present and past evolution than Darwin’s finches on the Galápagos Islands. Molecular phylogenetic studies have established relationships among the 18 species, the order and timing of branching, and the approximate age of the group of one million years (Lamichhaney et al. 2015, 2016; Chaves et al. 2016; Lawson and Petren 2017). The species diverged from each other in three key ecological traits: body size, and beak size and shape. Variation among species and individuals in these three traits reflects dietary specialization (Boag and Grant 1984; Grant 2017). Beak variation is subject to natural selection (Boag and Grant 1981) and influenced by introgressive hybridization (Grant and Grant 2014, 2020; Lamichhaney et al. 2020). As with human height, beak traits are highly heritable, and estimates of *h*^*2*^ for beak traits exceed 0.50 in three species, reaching 0.89 in *Geospiza fortis* (Keller et al. 2001; Grant and Grant 2014). Prior whole genome study of three ground-finch species (*Geospiza*) found that 28 loci are highly differentiated (Rubin et al. 2022). These loci represent ancestral haplotype blocks and contain genes significantly enriched for a role in craniofacial development. Importantly, it is not known whether these loci also control intraspecies variation.

We present the results of a whole-genome analysis of 3,958 birds from all four species of ground finches (*G. fortis, G. scandens, G. magnirostris*, and *G. fuliginosa*) on the small island of Daphne Major. We tracked individuals from these four species in their shared environment for over 30 years and measured their phenotypic evolution and fitness. We identify, quantify and document the importance of six large effect loci and show how allelic variation has changed under contrasting influences of natural selection and introgressive hybridization. According to these findings, a few loci of large effect contributed disproportionately to the rapid diversification of species in this classical example of adaptive radiation.

## Results

### Whole-genome community analysis over multiple generations

Blood samples were collected annually by B.R.G from the four species of finches on Daphne in the years 1988-2012 (from individuals hatched as early as 1983). We performed lowpass, whole-genome sequencing for all individuals captured. In total, we sequenced 3,958 individuals to a mean depth of 2.2x, comprising 1,913 *G. fortis*, 855 *G. scandens*, 582 *G. magnirostris*, 55 *G. fuliginosa*, and 553 individuals of hybrid origin (see Materials and Methods). For 2,543 individuals sampled as adults, P.R.G. measured three beak dimensions (length, depth, and width), body weight, and recorded sex when known. We employed an iterative imputation pipeline using a reference panel of 433 Darwin’s finches sequenced to higher coverage (15 ± 8X), a *de novo* pedigree-based recombination map (Fig. S1, Table S1), and the software GLIMPSE (Rubinacci et al. 2021), which imputes genotypes based on genotype likelihoods (*n*_*SNPs*_ = 5,163,840). We found high concordance in imputing variants with reference panel frequencies > 0.5% (Fig. S2) and concordance in genomic PCA using genotype-likelihoods alone (Fig. S3).

### Admixture and immigration are important components of population history

Genome-wide divergence is low among the four *Geospiza* species present on Daphne (*F*_ST_ = 0.03-0.17). To track the ancestry of every individual on the island, we estimated genomic ancestry using a set of ancestry informative and putatively neutral, unlinked markers (Methods). Daphne finches show extensive signatures of admixture, consistent with previous genomic and pedigree-based observations (Grant et al. 2004; Grant and Grant 2010, 2020; Lamichhaney et al. 2020). Reflecting the accumulated effect of introgression, finch species tended to be more genetically differentiated at the beginning than at the end of the study period; self-ancestry declined 20% in *G. fortis*, 17% in *G. scandens*, and 0.2% in *G. magnirostris* (Fig. 1). These are likely the lower bounds of ancestry due to high allele sharing across the radiation (Lamichhaney et al. 2015). The largest introgression of genomic material in *G. scandens* is from *G. fortis*, which increased by 12%. In contrast, the largest contribution to the *G. fortis* population originates from *G. fuliginosa*, whose ancestry increased 14%. These changes are consistent with pedigree information (Grant and Grant 2020) and are in stark contrast to the stability of *G. magnirostris* ancestry across the period (Fig. 1C).

**Fig. 1:**
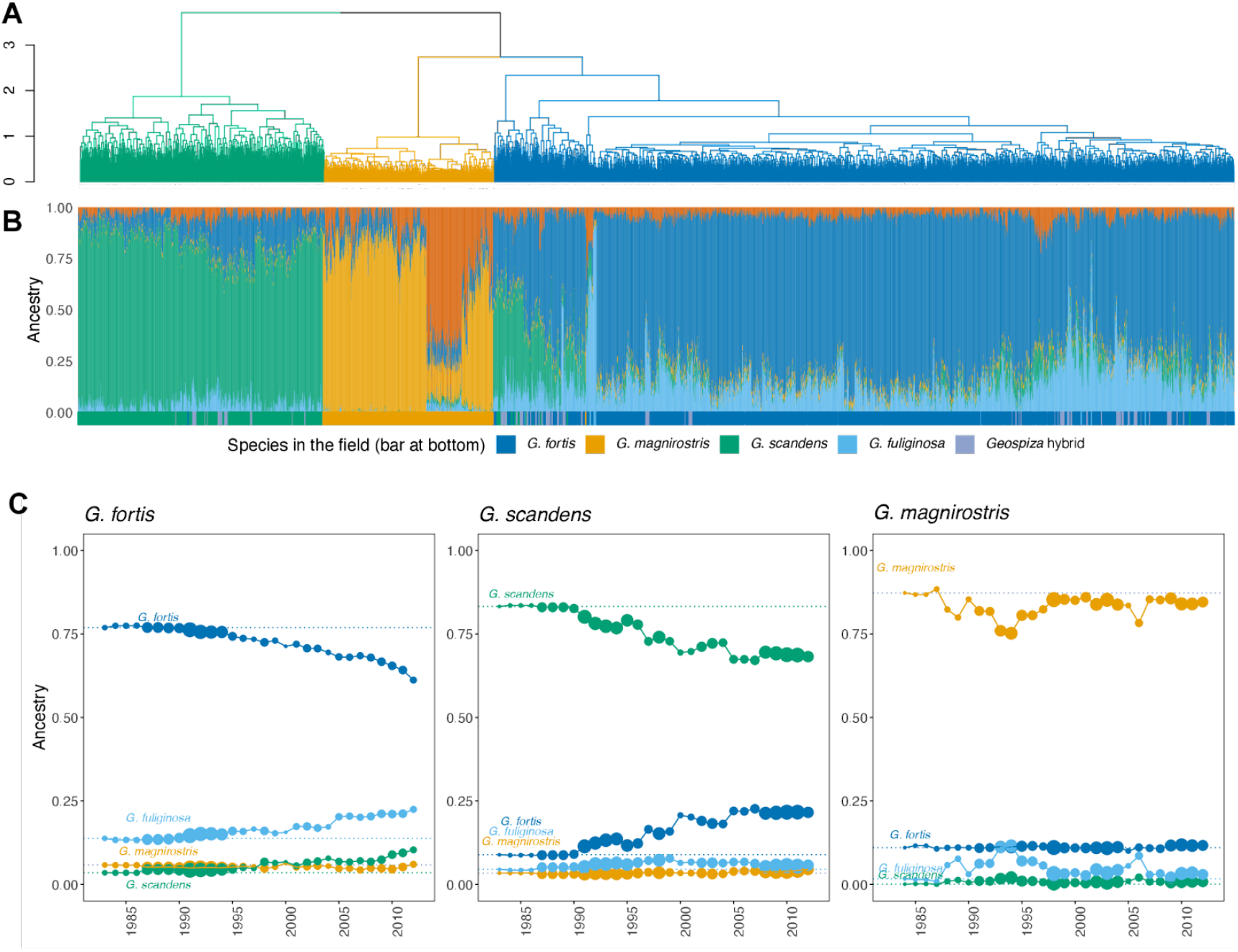
Admixture history of four species of finches on Daphne Major. (A) Clustering derived from the relatedness matrix produced using genome-wide SNPs in the software GEMMA. (B) Ancestry estimates (*K = 5*) for each of 3,958 individuals on Daphne based on ancestry informative SNPs (see Methods). Filled column colors designate the ancestral population containing a majority of samples from the field identification for each species by their measurements. Darker orange indicates a second *G. magnirostris* population. Below, a filled horizontal bar designates the field identification. (C) Annual trends in ancestry per species grouping from *K* = 4. Each panel refers to the field identification labeled above. Ancestry estimates are the mean value per cohort each year starting in 1983 and ending in 2012.

Changes in ancestry are strongly correlated with phenotypic shifts. Variation in beak morphology among the four Daphne finch species can be decomposed into two principal components explaining 99.7% of morphological variation, with PC1 loading on beak size (89% of variation, beak size hereinafter) and PC2 on beak shape (10% of variation, beak shape hereinafter, Fig. 2A). In *G. fortis*, change in beak shape is associated with *G. scandens* ancestry (*R*^2^ = 0.23, Fig. S5) and beak size with *G. fuliginosa* ancestry (*R*^2^ = 0.12, Fig. S5). In *G. scandens, G. fortis* ancestry is associated with beak shape (*R*^2^ = 0.34) and *G. fortis* and *G. fuliginosa* both contribute to beak size, but to a small extent (*R*^2^ = 0.04 and *R*^2^ = 0.01, Fig. S5). *G. scandens* and *G. fuliginosa* do not hybridize on Daphne, yet alleles pass from *G. fuliginosa* to *G. scandens* and these associations are the outcome of introgression through *G. fortis* as a conduit species (Supplemental Text 1, Fig. S4, also (Grant and Grant 2020).

**Fig. 2:**
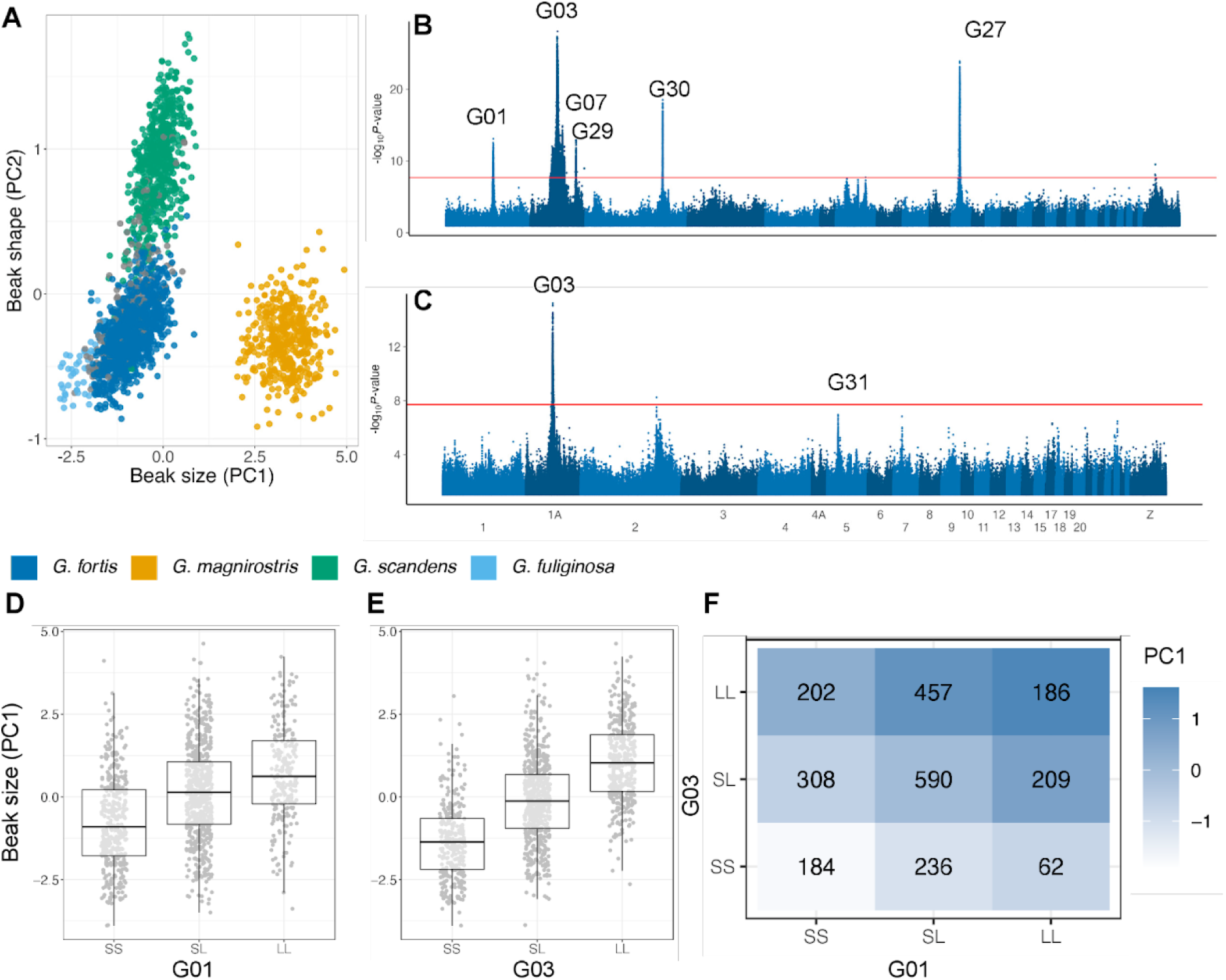
Genome-wide association analysis of morphological variation in beak and body size in *G. fortis*. (A) Morphological PCA for beak width, length, and depth, colored by species. (B) Multivariate genome-wide association analysis for beak PC1 and PC2, including body weight and sex as covariates. The cutoff for genome-wide significance at -log10(*P-*value) = 7.7 is indicated. Locus names match and extend those previously reported (Rubin et al. 2022). (C) Genome-wide association analysis for body weight using sex as a covariate. *G03* (*HMGA2*) and *G31*, containing *IGF2*, are highlighted. (D-E) Relationships between *G01* and *G03* genotypes and beak size. (F) Phenotypic effects of all genotype combinations at *G01* and *G03* suggest an additive relationship.

The estimated number of ancestral populations on Daphne is a good fit to *K* = 4 or to *K =* 5, i.e., one more than the expected number of species (Fig. S6). At *K* = 5, the fifth population reflects heterogeneity in the Daphne *G. magnirostris* population. This population was initiated by a single pair of individuals but augmented by immigrants later in the study period (Grant et al. 2001). Our ancestry analysis identifies two sources of immigrants to Daphne, one from the nearby island of Santa Cruz and a second that is ∼5% larger from an as-yet unknown island of origin (Fig. S7, Supplemental Text 2). These results highlight the dual contributions of hybridization and immigration to genetic and phenotypic diversity on Daphne.

### High SNP heritability of beak morphology and body size

We used the extensive genotype-phenotype records to determine SNP heritability of beak morphology and body size. Estimates of SNP heritability (*h*^*2*^ _SNP_) account for the proportion of variance in phenotypic traits explained by our imputed SNP dataset. Here we analyzed three phenotypic traits: beak size, beak shape, and body size using individuals with complete phenotypic data (n = 2,545). We ran all association analyses separately on three genetic clusters (Fig. 1A) representative of *G. fortis* (n =1,508), *G. scandens* (n = 552), and *G. magnirostris* (n = 430). We did not include *G. fuliginosa* samples in this analysis due to low power for genotype-phenotype analysis (*n* = 55).

To estimate *h*^*2*^_SNP_ we used a linkage disequilibrium (LD) and minor allele frequency (MAF)-stratified residual maximum likelihood analysis (GREML-LDMS) as implemented in the software package GCTA (Yang et al. 2015). We estimated a total beak size *h*^*2*^ _SNP_ in *G. fortis* of 0.95 (se = 0.02). A large proportion of this estimate is captured by common (MAF > 0.05) and high LD variants (*h*^*2*^_SNP_ = 0.77, se = 0.04, Fig. S8). Heritabilities of beak shape (*h*^*2*^ = 0.78, se = 0.03, Fig. S8) and weight (*h*^*2*^_SNP_ = 0.67, se = 0.04, Fig. S8) are also high (Keller et al. 2001; Grant and Grant 2014). The high SNP-based estimates match pedigree *h*^*2*^_SNP_ (Keller et al. 2001; Grant and Grant 2014) and confirm that beak traits and body size are highly heritable in Darwin’s finches.

### GWAS identifies large effect loci underlying ecological traits

To identify loci underlying phenotypic variation we performed GWAS using the software GEMMA (Zhou and Stephens 2014). Because beak and body size (weight) are strongly correlated (*G. fortis, r* = 0.74, *P* < 0.001), we included body weight as a covariate in a multivariate GWAS of beak size (PC1) and shape (PC2) as the response variables. For beak size and shape in *G. fortis*, we identified six independent loci surpassing a significance threshold of -log_10_(*P*) > 7.7 set by permutation (Supplemental Text 3, Fig. 2B, Fig. 3A-D, Fig. S9). A large region of association on chromosome 1A contains the previously identified *G03* locus encompassing *HMGA2* (Fig. 3B)(Lamichhaney et al. 2016). We treat this large region of association as a single locus following exploratory analysis that indicates long-range linkage disequilibrium extending from the central region of association (Fig. S10).

**Fig. 3:**
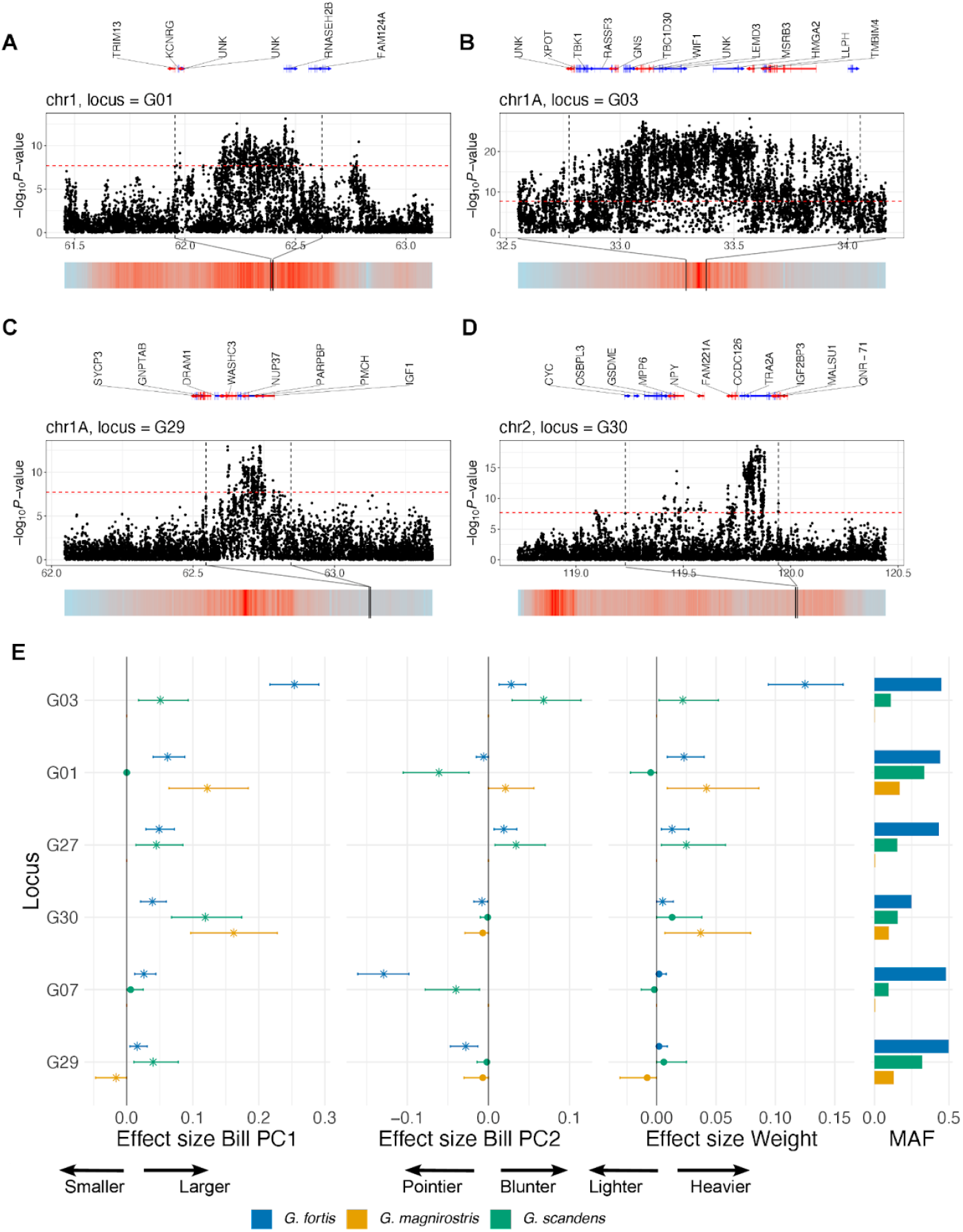
Details of association peaks and effect size estimates. A-D zoom-in of regions of associations for locus *G01, G03, G29*, and *G30*. Red points indicate missense mutations. Below, a heatmap of a sliding window of linkage disequilibrium of all SNPs in 200kb windows (blue = low, red = high). (E) Additive effect size predictions for each of the six loci shown in Fig. 2C. Colors indicate the three species, and a star designates statistical significance (*P*<0.05; see Methods). Right, the minor allele frequency for each locus across species. Note that *G. magnirostris* is fixed for the large allele at *G03, G07*, and *G29*.

We analyzed body weight using the relatedness matrix and sex as covariates. In *G. fortis*, we identified a large effect locus *G03* on chromosome 1A and two additional loci, *G30* and *G31*, that approached genome-wide significance (Fig. 2C). Thus, *G03* has a large effect on body weight and on beak size independent of body weight. These six loci also reach statistical significance in *G. scandens*, but only one (*G01*) does so in *G. magnirostris* (Fig. S11 and Fig. S12). Weaker associations in these species may reflect the smaller sample sizes (3x more samples in *G. fortis*) and less within-group variation at these loci. For example, *G. magnirostris* is nearly fixed for one allele at both *G03* and *G30* (and therefore GWAS does not detect these in this species, Fig. 2E, Fig. S12). A striking feature of *G. scandens* is an association of a single large region (34Mbp) on chromosome 5 (Fig. S12) that is associated with large divergent haplotypes (Fig. S13) and overlaps a region known to be resistant to ingression in *G. scandens* (Lamichhaney et al. 2020).

### How individual loci contribute to beak variation within populations

To estimate effect sizes, we identified haplotypes associated with each GWAS signal (Materials and Methods). In combination, the six loci account for 46% of the variation in beak size of *G. fortis*, and 19% of the variation in beak shape (Fig. 3E). These values reduce to 29% and 20%, respectively, after controlling for the correlated effects of body size by using residuals of a model including body weight and sex as a covariate (Fig. S15). Similarly, in *G. scandens*, these six loci account for 29% of beak size (residuals = 26%) and 24% of beak shape (residuals = 21%). All loci contribute additively to beak size variation (Fig. 2D-F, Fig. S18). When we fitted these six loci as covariates in a GREML analysis of *G. fortis* we found that they explain 59% of total *h*^2^ _SNP_ in beak size, 17% of the variation in weight, but only 3% of beak shape (see also Fig. S16). Together, these results demonstrate that a small number of loci explain a large portion of the heritable beak size variation in *G. fortis*. Effect size predictions were largely robust to filtering based on genomic ancestry estimates, suggesting that their magnitude is not inflated by background ancestry (Fig. S17).

The single largest contributor to beak size is *G03* which accounts for 23% of beak size variation in *G. fortis* (Fig. 3E), half of the total explained variation (46%). This is reduced to 11% after controlling for the correlated effect of body size. The remaining five loci explain between 2% (*G29*) and 7% (*G01*) of beak size variation. After controlling for body size, approximately 13% of beak shape variation is explained by another locus (*G07*). The remaining loci associated with beak size explain between 0% (*G30*) and 3% (*G29*) of variation in beak shape. At four loci, we detected a third haplotype in *G. scandens* associated with beak morphology (Supplemental Text 5, Table S2).

### Inter and intraspecies variation

Species of Darwin’s finches differ in beak size and shape; therefore, loci that explain differences between species may contribute to variation within species. Among 28 previously identified loci with large allele frequency differences among *Geospiza* species (Rubin et al. 2022), we identified four significant associations in our GWAS analysis (*G01, G03, G07*, and *G27*).

Are the remaining inter-species differences associated with intra-population variation? After pruning for LD-linked loci (Fig. S19), 6 out of the 10 remaining loci from Rubin et al. (2022) either had small effects on beak size (4 explained <1% of variation, Fig. S20) or explained 1-2% of variation in beak shape (2 loci) Fig. S20). In all 6 cases, the allele associated with a large beak in the *G. fortis* population on Daphne is the most common in the largest species, *G. magnirostris*, rare in the smallest species *G. fuliginosa*, and at intermediate frequency in *G. fortis* (Fig. S20). These include *G26*, an association overlapping *IGFBP2*, encoding an insulin growth factor binding proteins. Together these results indicate that species differences in allele frequency and beak and body size are to a large extent recapitulated in individual variation in *G. fortis*. Four loci with allele frequency variation among species on chromosome 2 were not associated with either beak size or shape in *G. fortis* and may therefore affect other fitness traits.

### Two novel loci involved in beak size

Two loci in this paper have not previously been associated with individual variation in Darwin’s finches (*G29* and *G30*) and each contain insulin-like growth factor-related genes (*IGF1* and *IGF2BP3*, Fig. 3C and D); genes that are part of a well-characterized network involved in growth, metabolism, and aging (Wullschleger et al. 2006; Zoncu et al. 2011; Melzer et al. 2020; Morrill et al. 2022). Insulin-like growth factors are also associated with beak size in *Pyrenestes ostrinus* (vonHoldt et al. 2018) and the evolution of life history variation across amniotes (McGaugh et al. 2015). A third IGF-related gene, *IGF2*, falls just short of genome-wide significance in the body size GWAS in *G. fortis* (*G31*). Together with *G26 (IGFBP2)*, the identification of four *IGF* loci involved in Darwin’s finch beak morphology supports the hypothesis that gene networks involved in this pathway are co-evolving (McGaugh et al. 2015).

### How natural selection and hybridization change allele frequencies at major effect loci

Populations of *G. fortis* and *G. scandens* underwent substantial change in beak morphology over the 30 years of monitoring (Fig. 4A) (Grant and Grant 2014). The average size of *G. fortis* and *G. scandens* beak dimensions oscillated in direction in response to changing ecological conditions, with an abrupt change to smaller beaks in *G. fortis* and a more gradual change to blunter beaks in *G. scandens* (Fig. 4A) (Grant and Grant 2002). Allele frequency changes at the large-effect loci identified in our GWAS changed concordantly and exceeded annual allele frequency shifts at random loci (Fig. 4B, Fig. S21). The strongest shift occurred in *G. fortis* during the 2004/2005 drought (Grant and Grant 2006). The beaks of *G. fortis* became smaller on average due to differential mortality resulting from competition with the much larger *G. magnirostris*. We confirm the sharp increase in the frequency of the small *G03* haplotype (Fig. 4C) from 0.50 in 2004 to 0.63 in 2005 (Lamichhaney et al. 2016). *G03* predicts survival alone with a selection coefficient of 0.49 (Fisher’s exact test, *P* < 0.05). However, three other loci have differential effects in the same direction when comparing small to large homozygotes (and two others trend in this direction). In fact three loci (*G01, G03*, and *G29*, Fig. 4D) predict survival from 2004 to 2005 better than *G03* alone in a repeated leave-one-out analysis (AIC_*combined*_ = 89.4 vs AIC_*G03*_ = 97.6) (Fig. 4D, *z* = 3.1, *P* < 0.01).

**Fig. 4:**
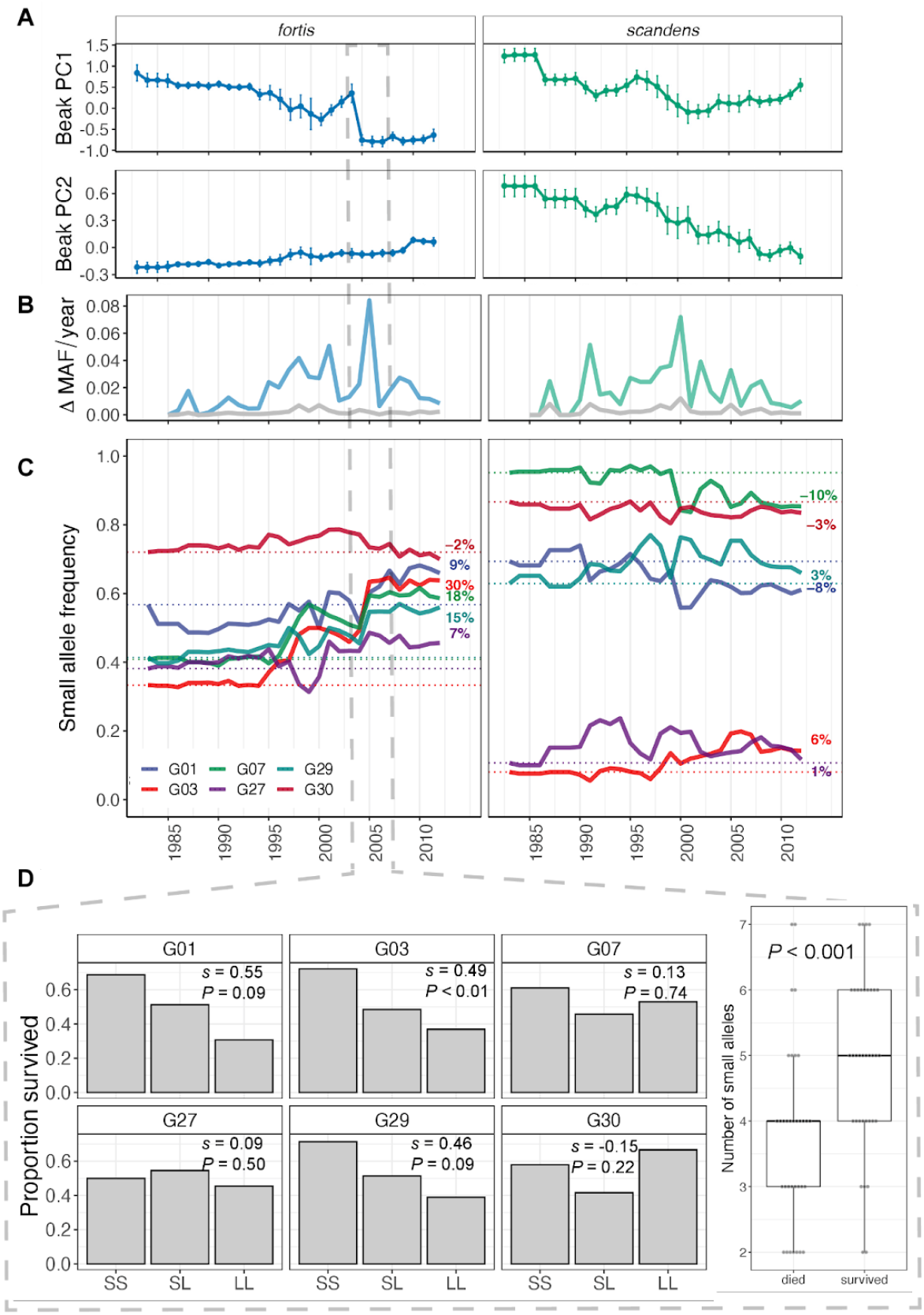
Evolutionary change over 30 years on Daphne Major. (A) Annual phenotypic means for *G. fortis* and *G. scandens* between 1983 and 2012. (B) The absolute average change in allele frequency (AF_year n_ - AF_year n-1_) across all six loci shown in Fig. 2. The colored line indicates the average of all six loci and the gray line indicates 100 randomly selected loci across the genomes with starting allele frequencies matching the six loci. (C) Annual allele frequency trajectories at each of the six loci. (D) The proportion of individuals that survived during the 2004/2005 drought at each of the six loci (*n =* 73 individuals). The selection coefficient for the large allele is indicated, and a Fisher’s exact test *P*-value comparing allele frequency before and after the drought is given (see Materials and Methods). Right, the sum of small alleles at *GF01, GF03*, and *GF29* is associated with survival during the 2004/2005 drought event.

How has changing ancestries (Fig. 1) affected allele frequency change at these six loci? Allele frequency changes at these six loci range from 2-30% in *G. fortis* and from 1-10% in *G. scandens* (Fig. 4C). The phenotypic change towards a blunter beak in *G. scandens* is the consequence of incremental gene flow from *G. fortis* beginning in the 1990s (Fig. 1) (Grant and Grant 2002; Lamichhaney et al. 2020). A spike in allele frequency change at the turn of the century is likely due to this introgression, which occurred following the low productivity of *G. scandens* in the El Niño year of 1998 (Fig. 1) (Grant and Grant 2014). At the beak shape locus with the largest phenotypic effect (*G07/ALX1*), *G. scandens* carrying a blunt allele are more likely to have *G. fortis* ancestry than those carrying a pointed allele (GLM: *F*=1.1, *P* < 0.0001, see also Supplemental Text 1). Similarly, *G. fortis* carrying the small allele at *G03*, the locus of greatest phenotypic effect on beak size, are more likely to have *G. fuliginosa* ancestry (GLM: *F*=-1.1, *P* < 0.0001).

## Conclusions

We describe the genetic architecture underlying phenotypic variation and fitness in natural populations of Darwin’s finches using lowpass, whole-genome sequencing over 5-6 generations on the undisturbed island of Daphne Major, Galápagos. We report that six large effect loci are responsible for as much as 46% of the genetic variance in beak size in *G. fortis* and that one alone (*G03/HMGA2*) explains 23% of the variance. The latter is due to a QTL effect on allometric growth (affecting both body and beak size) and an additional QTL effect on beak size independent of body size.

These results represent a striking deviation from a solely polygenic model involving many loci of small effect assumed to underlie most phenotypic traits (Rockman 2012; Barton 2022; Campagna and Toews 2022). However, in wild populations, an increasing number of studies show that large effect loci (Shapiro et al. 2004; Chan et al. 2010; Barrett et al. 2019; Han et al. 2020; Schluter and Rieseberg 2022) play a more important role in maintaining phenotypic variation in natural populations than anticipated from GWAS in humans (Manolio et al. 2009) and artificial selection data (Goddard et al. 2016). Why do these discrepancies in genetic architecture occur? An important difference is that most human genotype-phenotype relationships explored by GWAS depend on standing genetic variation, including slightly deleterious variants not yet purged by natural selection (Zeng et al. 2018). Further, responses to artificial selection reflect short-term directional selection on standing variation. In contrast, the data on Darwin’s finches and other natural populations deviating from a solely polygenic model represent long-term responses to intense perturbations by natural selection, introgression, and environmental fluctuations. In humans, large effect alleles are known to affect adaptation to extreme environments at high altitudes (Simonson et al. 2010) and diving (Ilardo et al. 2018), and following strong selection associated with the Black Death (Klunk et al. 2022).

Genetic theory predicts that large effect loci should be quickly fixed by selection (Barton 2022). However, that is only possible when conditions favoring a positive effect on fitness remain until fixation has occurred. For example, the small and large alleles at the six loci described here are close to fixation in the small (*G. fuliginosa*) and large ground-finches (*G. magnirostris*), respectively. Conditions that vary over time may instead lead to balancing selection where the fitness of alleles fluctuates in time or varies in space. This is the case in the Galápagos, where oscillating environmental conditions have unpredictable evolutionary outcomes (Grant and Grant 2002). We found substantial shifts in allele frequencies at these large effect loci under periods of environmental stress and intense interspecies competition in *G. fortis*, which has an intermediate phenotype (Fig. 4). Another important factor promoting segregation of large effect alleles in Darwin’s finches, and in other species (Seehausen et al. 2014; Edelman et al. 2019), is gene flow between closely related populations. We show that introgression supplements standing variation at large effect loci in *G. fortis*, leading to intermediate genotype frequencies (Fig. 3, bottom) that facilitate survival under the unpredictable environmental conditions of Daphne (Grant and Grant 2002). These results are consistent with theory predicting that adaptation combined with migration favors the evolution of large effect alleles (Yeaman and Whitlock 2011).

Large effect mutations under positive selection are rare. Therefore, a more plausible interpretation is that the large effect alleles detected in the present study represent haplotypes harboring multiple mutations affecting fitness. Such large effect alleles have evolved in domestic animals by the consecutive accumulation of multiple mutations (Marklund et al. 1998; Andersson and Purugganan 2022). A similar evolution of large effect alleles in natural populations has also been noted (Prud’homme et al. 2006; McGregor et al. 2007). Thus, the six large effect loci detected in this study may be considered small supergenes where the haplotype blocks are maintained because they are located in low-recombination regions and recombinant haplotypes are eliminated by selection. The two major *G03/HMGA2* alleles represent ∼525 kb ancient haplotypes, encompassing four genes - *HMGA2, MSRB3, LEMD3, WIF1*- and most likely have affected variation in body size and beak morphology throughout the evolutionary history of Darwin’s finches (Rubin et al. 2022; Lamichhaney et al. 2016). Given the fundamental role for *HMGA2* as a transcription-facilitating factor controlling body size in vertebrates (Lee et al. 2022), we hypothesize that non-coding changes affecting *HMGA2* expression regulate development of body and beak size. However, the additional effect of this locus on beak size independent of body size found here may be controlled by mutation(s) affecting the function of one or more of the other genes at this locus.

Taken together, this community-level genomic study of Darwin’s finch populations on the small island of Daphne has demonstrated a prominent role of large effect alleles at a few loci in causing evolutionary change when the populations are subjected to intense natural selection and influenced by introgressive hybridization.

## Acknowledgments

We thank R. Corbett-Detig for insightful discussion, A. Cocco and J. Pettersson for assistance in the lab. The National Genomics Infrastructure (NGI)/Uppsala Genome Center provided service in massive parallel sequencing, and the computational infrastructure was provided by the Swedish National Infrastructure for Computing (SNIC) at UPPMAX, partially funded by the Swedish Research Council through grant agreement no. 2018-05973. NSF (USA) funded the collection of material under permits from the Galápagos and Costa Rica National Parks Services and the Charles Darwin Research Station, and in accordance with protocols of Princeton University’s Animal Welfare Committee.

## Funding

The project was financially supported by Vetenskapsrådet (2017-02907) and Knut and Alice Wallenberg Foundation (KAW 2016.0361).

## Author contributions

Conceptualization: LA, PRG, BRG, EDE Data Curation: CGS, EDE

Methodology: EDE, LA

Formal analysis: EDE, ASP, CJR, PV Investigation: CGS, EDE, PRG, BRG Visualization: EDE, ASP

Funding acquisition: LA, PRG, BRG Resources: PRG, BRG

Supervision: LA

Writing – original draft: EDE

Writing – review & editing: all authors

## Competing interests

Authors declare that they have no competing interests.

## Supplementary Materials

Materials and Methods

Supplementary Text S1 to S5

Figs. S1 to S21

Tables S1 to S2

References (##–##)

## Notes

### Competing Interest Statement

The authors have declared no competing interest.

## References

Andersson, L., and M. Purugganan. 2022. Molecular genetic variation of animals and plants under domestication. Proceedings of the National Academy of Sciencesx 119:e2122150119.

Barrett, R. D. H., S. Laurent, R. Mallarino, S. P. Pfeifer, C. C. Y. Xu, M. Foll, K. Wakamatsu, J. S. Duke-Cohan, J. D. Jensen, and H. E. Hoekstra. 2019. Linking a mutation to survival in wild mice. Science 363:499–504.

Barton, N. H. 2022. The “New Synthesis.” Proceedings of the National Academy of Sciences 119:e2122147119.

Boag, P. T., and P. R. Grant. 1981. Intense natural selection in a population of Darwin’s finches (geospizinae) in the Galápagos. Science 214:82–85.

Boag, P. T., and P. R. Grant. 1984. The classical case of character release: Darwin’s finches (Geospiza) on Isla Daphne Major, Galápagos. Biol. J. Linn. Soc. Lond. 22:243–287. Oxford University Press (OUP).

Bosse, M., L. G. Spurgin, V. N. Laine, E. F. Cole, J. A. Firth, P. Gienapp, A. G. Gosler, K. McMahon, J. Poissant, I. Verhagen, M. A. M. Groenen, K. van Oers, B. C. Sheldon, M. E. Visser, and J. Slate. 2017. Recent natural selection causes adaptive evolution of an avian polygenic trait. Science 358:365–368.

Campagna, L., M. Repenning, L. F. Silveira, C. S. Fontana, P. L. Tubaro, and I. J. Lovette. 2017. Repeated divergent selection on pigmentation genes in a rapid finch radiation. Science Advances 3:e1602404.

Campagna, L., and D. P. L. Toews. 2022. The genomics of adaptation in birds. Curr. Biol. 32:R1173–R1186. Elsevier.

Chan, Y. F., M. E. Marks, F. C. Jones, G. Villarreal Jr, M. D. Shapiro, S. D. Brady, A. M. Southwick, D. M. Absher, J. Grimwood, J. Schmutz, R. M. Myers, D. Petrov, B. Jónsson, D. Schluter, M. A. Bell, and D. M. Kingsley. 2010. Adaptive evolution of pelvic reduction in sticklebacks by recurrent deletion of a Pitx1 enhancer. Science 327:302–305.

Chaves, J. A., E. A. Cooper, A. P. Hendry, J. Podos, L. F. D. León, J. A. M. Raeymaekers, O. W. McMillan, and J. A. C. Uy. 2016. Genomic variation at the tips of the adaptive radiation of Darwin’s finches. Mol. Ecol., doi: 10.1111/mec.13743.

Edelman, N. B., P. B. Frandsen, M. Miyagi, B. Clavijo, J. Davey, R. B. Dikow, G. García-Accinelli, S. M. Van Belleghem, N. Patterson, D. E. Neafsey, R. Challis, S. Kumar, G. R. P. Moreira, C. Salazar, M. Chouteau, B. A. Counterman, R. Papa, M. Blaxter, R. D. Reed, K. K. Dasmahapatra, M. Kronforst, M. Joron, C. D. Jiggins, W. O. McMillan, F. Di Palma, A. J. Blumberg, J. Wakeley, D. Jaffe, and J. Mallet. 2019. Genomic architecture and introgression shape a butterfly radiation. Science 366:594–599.

Goddard, M. E., K. E. Kemper, I. M. MacLeod, A. J. Chamberlain, and B. J. Hayes. 2016. Genetics of complex traits: prediction of phenotype, identification of causal polymorphisms and genetic architecture. Proc. Biol. Sci. 283:1173–1186.

Grant, P. R. 2017. Ecology and Evolution of Darwin’s Finches (Princeton Science Library Edition). Princeton University Press.

Grant, P. R., and B. R. Grant. 2010. Conspecific versus heterospecific gene exchange between populations of Darwin’s finches. Philos. Trans. R. Soc. Lond. B Biol. Sci. 365:1065–1076.

Grant, P. R., and B. R. Grant. 2006. Evolution of character displacement in Darwin’s finches. Science 313:224–226.

Grant, P. R., and B. R. Grant. 2020. Triad hybridization via a conduit species. Proc. Natl. Acad. Sci. U. S. A. 117:7888–7896.

Grant, P. R., and B. R. Grant. 2002. Unpredictable evolution in a 30-year study of Darwin’s finches. Science 296:707–711.

Grant, P. R., B. R. Grant, J. A. Markert, L. F. Keller, and K. Petren. 2004. Convergent Evolution of Darwin’S Finches Caused By Introgressive Hybridization and Selection. Evolution 58:1588–1599.

Grant, P. R., B. R. Grant, and K. Petren. 2001. A population founded by a single pair of individuals: Establishment, expansion, and evolution. Genetica 112-113:359–382.

Grant, P. R., and R. B. Grant. 2014. 40 Years of Evolution: Darwin’s Finches on Daphne Major Island. Princeton University Press.

Han, F., M. Jamsandekar, M. E. Pettersson, L. Su, A. P. Fuentes-Pardo, B. W. Davis, D. Bekkevold, F. Berg, M. Casini, G. Dahle, E. D. Farrell, A. Folkvord, and L. Andersson. 2020. Ecological adaptation in Atlantic herring is associated with large shifts in allele frequencies at hundreds of loci. Elife 9.

Ilardo, M. A., I. Moltke, T. S. Korneliussen, J. Cheng, A. J. Stern, F. Racimo, P. de Barros Damgaard, M. Sikora, A. Seguin-Orlando, S. Rasmussen, I. C. L. van den Munckhof, R. Ter Horst, L. A. B. Joosten, M. G. Netea, S. Salingkat, R. Nielsen, and E. Willerslev.2018. Physiological and Genetic Adaptations to Diving in Sea Nomads. Cell 173:569–580.e15.

Keller, L. F., P. R. Grant, B. R. Grant, and K. Petren. 2001. Heritability of morphological traits in Darwin’s finches: misidentified paternity and maternal effects. Heredity 87:325–336.

Klunk, J., T. P. Vilgalys, C. E. Demeure, X. Cheng, M. Shiratori, J. Madej, R. Beau, D. Elli, M. I. Patino, R. Redfern, S. N. DeWitte, J. A. Gamble, J. L. Boldsen, A. Carmichael, N. Varlik, K. Eaton, J.-C. Grenier, G. B. Golding, A. Devault, J.-M. Rouillard, V. Yotova, R. Sindeaux, C. J. Ye, M. Bikaran, A. Dumaine, J. F. Brinkworth, D. Missiakas, G. A. Rouleau, M. Steinrücken, J. Pizarro-Cerdá, H. N. Poinar, and L. B. Barreiro. 2022. Evolution of immune genes is associated with the Black Death. Nature, doi: 10.1038/s41586-022-05349-x.

Lamichhaney, S., J. Berglund, M. S. Almén, K. Maqbool, M. Grabherr, A. Martinez-Barrio, M. Promerová, C.-J. Rubin, C. Wang, N. Zamani, B. R. Grant, P. R. Grant, M. T. Webster, and L. Andersson. 2015. Evolution of Darwin’s finches and their beaks revealed by genome sequencing. Nature 518:371–375.

Lamichhaney, S., F. Han, J. Berglund, C. Wang, M. S. Almén, M. T. Webster, B. R. Grant, P. R. Grant, and L. Andersson. 2016. A beak size locus in Darwin’s finches facilitated character displacement during a drought. Science 352:470–474.

Lamichhaney, S., F. Han, M. T. Webster, B. R. Grant, P. R. Grant, and L. Andersson. 2020. Female-biased gene flow between two species of Darwin’s finches. Nature Ecology & Evolution 4:979–986.

Lawson, L. P., and K. Petren. 2017. The adaptive genomic landscape of beak morphology in Darwin’s finches. Mol. Ecol. 26:4978–4989.

Lee, M. O., J. Li, B. W. Davis, S. Upadhyay, H. M. Al Muhisen, L. J. Suva, T. M. Clement, and L. Andersson. 2022. Hmga2 deficiency is associated with allometric growth retardation, infertility, and behavioral abnormalities in mice. G3 12.

Manolio, T. A., F. S. Collins, N. J. Cox, D. B. Goldstein, L. A. Hindorff, D. J. Hunter, M. I. McCarthy, E. M. Ramos, L. R. Cardon, A. Chakravarti, J. H. Cho, A. E. Guttmacher, A. Kong, L. Kruglyak, E. Mardis, C. N. Rotimi, M. Slatkin, D. Valle, A. S. Whittemore, M. Boehnke, A. G. Clark, E. E. Eichler, G. Gibson, J. L. Haines, T. F. C. Mackay, S. A. McCarroll, and P. M. Visscher. 2009. Finding the missing heritability of complex diseases. Nature 461:747–753.

Marklund, S., J. Kijas, H. Rodriguez-Martinez, L. Rönnstrand, K. Funa, M. Moller, D. Lange, I. Edfors-Lilja, and L. Andersson. 1998. Molecular basis for the dominant white phenotype in the domestic pig. Genome Res. 8:826–833.

Marques, D. A., J. I. Meier, and O. Seehausen. 2019. A Combinatorial View on Speciation and Adaptive Radiation. Trends Ecol. Evol. 34:531–544. Elsevier Ltd.

McGaugh, S. E., A. M. Bronikowski, C.-H. Kuo, D. M. Reding, E. A. Addis, L. E. Flagel, F. J. Janzen, and T. S. Schwartz. 2015. Rapid molecular evolution across amniotes of the IIS/TOR network. Proc. Natl. Acad. Sci. U. S. A. 112:7055–7060.

McGee, M. D., S. R. Borstein, J. I. Meier, D. A. Marques, S. Mwaiko, A. Taabu, M. A. Kishe, B. O’Meara, R. Bruggmann, L. Excoffier, and O. Seehausen. 2020. The ecological and genomic basis of explosive adaptive radiation. Nature 586:75–79. Springer US.

McGregor, A. P., V. Orgogozo, I. Delon, J. Zanet, D. G. Srinivasan, F. Payre, and D. L. Stern. 2007. Morphological evolution through multiple cis-regulatory mutations at a single gene. Nature 448:587–590.

Meier, J. I., D. A. Marques, C. E. Wagner, L. Excoffier, and O. Seehausen. 2018. Genomics of parallel ecological speciation in Lake Victoria cichlids. Mol. Biol. Evol. 35:1489–1506.

Melzer, D., L. C. Pilling, and L. Ferrucci. 2020. The genetics of human ageing. Nat. Rev. Genet. 21:88–101.

Morrill, K., J. Hekman, X. Li, J. McClure, B. Logan, L. Goodman, M. Gao, Y. Dong, M. Alonso, E. Carmichael, N. Snyder-Mackler, J. Alonso, H. J. Noh, J. Johnson, M. Koltookian, C. Lieu, K. Megquier, R. Swofford, J. Turner-Maier, M. E. White, Z. Weng, A. Colubri, D. P. Genereux, K. A. Lord, and E. K. Karlsson. 2022. Ancestry-inclusive dog genomics challenges popular breed stereotypes. Science 376:eabk0639.

Prud’homme, B., N. Gompel, A. Rokas, V. A. Kassner, T. M. Williams, S.-D. Yeh, J. R. True, and S. B. Carroll. 2006. Repeated morphological evolution through cis-regulatory changes in a pleiotropic gene. Nature 440:1050–1053.

Rockman, M. V. 2012. The QTN program and the alleles that matter for evolution: All that’s gold does not glitter. Evolution 66:1–17.

Rubinacci, S., D. M. Ribeiro, R. J. Hofmeister, and O. Delaneau. 2021. Efficient phasing and imputation of low-coverage sequencing data using large reference panels. Nat. Genet. 53:120–126.

Rubin, C.-J., E. D. Enbody, M. P. Dobreva, A. Abzhanov, B. W. Davis, S. Lamichhaney, M. Pettersson, A. T. Sendell-Price, C. G. Sprehn, C. A. Valle, K. Vasco, O. Wallerman, B. R. Grant, P. R. Grant, and L. Andersson. 2022. Rapid adaptive radiation of Darwin’s finches depends on ancestral genetic modules. Science Advances 8:eabm5982.

Schluter, D., and L. H. Rieseberg. 2022. Three problems in the genetics of speciation by selection. Proceedings of the National Academy of Sciences 119:e2122153119.

Seehausen, O., R. K. Butlin, I. Keller, C. E. Wagner, J. W. Boughman, P. A. Hohenlohe, C. L. Peichel, G.-P. Saetre, C. Bank, A. Brännström, A. Brelsford, C. S. Clarkson, F. Eroukhmanoff, J. L. Feder, M. C. Fischer, A. D. Foote, P. Franchini, C. D. Jiggins, F. C. Jones, A. K. Lindholm, K. Lucek, M. E. Maan, D. A. Marques, S. H. Martin, B. Matthews, J. I. Meier, M. Möst, M. W. Nachman, E. Nonaka, D. J. Rennison, J. Schwarzer, E. T. Watson, A. M. Westram, and A. Widmer. 2014. Genomics and the origin of species. Nat. Rev. Genet. 15:176–192.

Shapiro, M. D., M. E. Marks, C. L. Peichel, B. K. Blackman, K. S. Nereng, B. Jónsson, D. Schluter, and D. M. Kingsley. 2004. Genetic and developmental basis of evolutionary pelvic reduction in threespine sticklebacks. Nature 428:717–723.

Simonson, T. S., Y. Yang, C. D. Huff, H. Yun, G. Qin, D. J. Witherspoon, Z. Bai, F. R. Lorenzo, J. Xing, L. B. Jorde, J. T. Prchal, and R. Ge. 2010. Genetic evidence for high-altitude adaptation in Tibet. Science 329:72–75.

vonHoldt, B. M., R. Y. Kartzinel, C. D. Huber, V. Le Underwood, Y. Zhen, K. Ruegg, K. E. Lohmueller, and T. B. Smith. 2018. Growth factor gene IGF1 is associated with bill size in the black-bellied seedcracker Pyrenestes ostrinus. Nat. Commun. 9:4855. Springer US.

Wullschleger, S., R. Loewith, and M. N. Hall. 2006. TOR signaling in growth and metabolism. Cell 124:471–484.

Yang, J., A. Bakshi, Z. Zhu, G. Hemani, A. A. E. Vinkhuyzen, S. H. Lee, M. R. Robinson, J. R. B. Perry, I. M. Nolte, J. V. van Vliet-Ostaptchouk, H. Snieder, LifeLines Cohort Study, T. Esko, L. Milani, R. Mägi, A. Metspalu, A. Hamsten, P. K. E. Magnusson, N. L. Pedersen, E. Ingelsson, N. Soranzo, M. C. Keller, N. R. Wray, M. E. Goddard, and P. M. Visscher. 2015. Genetic variance estimation with imputed variants finds negligible missing heritability for human height and body mass index. Nat. Genet. 47:1114–1120.

Yeaman, S., and M. C. Whitlock. 2011. The genetic architecture of adaptation under migration-selection balance. Evolution 65:1897–1911.

Zeng, J., R. de Vlaming, Y. Wu, M. R. Robinson, L. R. Lloyd-Jones, L. Yengo, C. X. Yap, A. Xue, J. Sidorenko, A. F. McRae, J. E. Powell, G. W. Montgomery, A. Metspalu, T. Esko, G. Gibson, N. R. Wray, P. M. Visscher, and J. Yang. 2018. Signatures of negative selection in the genetic architecture of human complex traits. Nat. Genet. 50:746–753.

Zhou, X., and M. Stephens. 2014. Efficient multivariate linear mixed model algorithms for genome-wide association studies. Nat. Methods 11:407–409.

Zoncu, R., A. Efeyan, and D. M. Sabatini. 2011. mTOR: from growth signal integration to cancer, diabetes and ageing. Nat. Rev. Mol. Cell Biol. 12:21–35.

