## Supplemental materials for "Large effect loci have a prominent role in Darwin’s finch evolution"

**This PDF file includes:**

Materials and Methods
Supplementary Text S1 to S5
Figs. S1 to S20
Tables S1 to S2
Captions for Data S1 to Sx

**Other Supplementary Materials for this manuscript include the following:**

Data S1 to Sx [paste data table titles in a list]

### Materials and Methods

#### *Sample collection*

Blood samples from finches on Daphne and various Galápagos islands were collected using sample protocols described elsewhere (Grant and Grant 2014; Lamichhaney et al. 2015, 2016). Briefly, blood was collected from the brachial vein, preserved on *EDTA*-soaked filter paper, and stored in Drierite. This includes the 3,358 samples on Daphne used in this study. Among these samples are 607 samples of both *G. fortis* and *G. scandens* samples used in a recent publication (Enbody et al. 2021). We also used a sequencing dataset of previously collected samples of all 18 species collected across the archipelago, Cocos Island, and two outgroup species from Barbados (Lamichhaney et al. 2015, 2016; Rubin et al. 2021). These samples were previously sequenced at moderate sequencing depth (15 ± 8 X). In addition, we sequenced 92 additional samples at moderate sequencing coverage (see below) for our reference panel. These were collected previously and represent 11 species from 14 islands.

#### *Field species identification*

Initial species assignment was determined based on measurements in the field, and we refer to a category of species identification as "Species in the field." This field identification is separate from the assignment of individuals to relatedness grouping in the GWAS analysis and from the assignment of genomic ancestry using Admixture (both described below). The field identification was used to direct analyses and at times, omit individuals from analysis (see the removal of *G. fuliginosa* individuals from GWAS analysis).

Identifying individuals to species followed early taxonomic descriptions of the species (Lack 1945) using beak morphology determined from measurements in the field as an initial identification. Identifying *G. fuliginosa* from small *G. fortis* is challenging because the distributions of beak measurements in the two species overlap. We used a classification scheme to identify *G. fuliginosa* immigrants based on measurements of finches from the northern coast of nearby Santa Cruz Island where the distributions of *G. fortis* and *G. fuliginosa* beak measurements do not overlap. Later in the study, *G. scandens* and *G. fortis* became increasingly morphologically similar following hybridization. Questionable individuals with intermediate morphology were assigned to the three species or to hybrids with alleles at 14 microsatellite loci. Individuals were classified as of hybrid origin if they were the offspring of a mixed species pairings detected during nest monitoring. See (Grant and Grant 2014) for a complete description of these methods.

#### *DNA extraction and library preparation*

We extracted DNA from blood on filter paper using a previously described custom salt preparation protocol (Enbody et al. 2021). We generated short-fragment libraries for whole-genome sequencing using a custom Tn5 transposon-based protocol that is available from https://www.protocols.io (Sprehn et al. 2021) and was described previously (Picelli et al. 2014; Enbody et al. 2021). Briefly, we targeted an insert size of 350bp for all samples and pooled individually barcoded samples for sequencing on a NovaSeq S4 flow cell (Illumina, CA) with a target depth of 2x. This was equivalent to roughly 400 samples per flow cell based on the ~1.1 Gbp genome size. To generate genotype calls, we first built a reference panel using higher coverage sequenced individuals from across the Galápagos islands (Lamichhaney et al. 2015, 2016; Rubin et al. 2022).

#### *Reference panel generation*

We used 321 previously published high-coverage samples of all 18 species of Darwin's finches to generate a first round of phased haplotype data to use as a reference panel. We mapped short-reads for these 321 individuals to the Camarhynchus_parvulus_V1.1 genome assembly (GCA_902806625.1) using the Sentieon tools wrapper (Freed et al. 2017) and the included BWA-mem v0.7.17 module (Li 2013). We called variants using a Sentieon tools version of Haplotype Caller (GATK 4.1) (Poplin et al. 2017) and joint genotyped all samples using Sentieon's GVCFtyper module (a wrapper around GenotypeGVCFs from GATK 4.1). Finally, we used using VariantFiltration (GATK v4.1.4.1) to apply the following filters to SNPs in the callset:
QD < 2.0, FS > 60.0, MQ < 40.0, MQRankSum < -12.5, ReadPosRankSum < -8.0, SOR > 3.0

In addition to genotype filters for:
DP < 2, DP > 100, GQ < 20

These filtered genotype calls were set to no call using Select Variants (GATK, v4.1.4.1) then removed all sites not passing filters using bcftools (http://www.htslib.org/, v1.10). We retained only biallelic SNPs for genotype phasing from this variant call set.

We statistically inferred haplotypes using a combination of WhatsHap (Martin et al. 2016) (v0.18) and SHAPEIT4 (Delaneau et al. 2019) (v4.1.3). First, we ran WhatsHap with default settings to identify phase-informative reads for each sample (i.e., a read spanning at least two heterozygous sites). Next, all individual VCFs with phase annotations were merged using the bcftools merge command. SHAPEIT4 was run with a constant recombination rate of 1 cM per

Mb and an expected error rate in the phase informative reads of 0.0001 (the default setting).

#### *Genotype imputation using GLIMPSE*

We used the SHAPEIT4 phased VCF as a reference panel to impute all low-coverage sequenced samples based on genotype likelihoods. We conducted imputation in two phases. We mapped reads to the Camarhynchus_parvulus_V1.1 genome assembly (GCA_902806625.1) as above using the BWA-mem module of Sentieon tools. We next generated a list of variant positions available in the reference panel by passing this file through `bcftools query `-f'%CHROM\t%POS\t%REF,%ALT\n' `. This file was used in a bcftools mpileup call for each sample to only calculate genotype likelihoods for known SNP positions. The bcftools (v1.10) command is as follows (as recommended in the GLIMPSE documentation):

```
bcftools mpileup --threads 2 -f ${REFGEN} -I -E -a 'FORMAT/DP' -T ${VCF}
${BAM} -Ou | bcftools call -Aim -C alleles -T ${TSV} -Oz -o ${OUT}
```

For the first phase imputation, we used the imputation software GLIMPSE (Rubinacci et al. 2021)(V1.1) to impute genotypes without a genetic map. GLIMPSE is designed for low-coverage sequence projects and uses genotype-likelihoods, rather than genotype calls, to impute genotypes from a reference panel. GLIMPSE is run in four stages: First, we divided the reference panel into 2Mb chunks with a buffer size of 200kb for efficient parallelization using GLIMPSE_chunk. Next, we ran the main algorithm, GLIMPSE_phase (without passing a genetic map), for 15 iterations to impute genotypes. Chunks were then ligated together using GLIMPSE_ligate and genotypes were output using GLIMPSE_sample and the –solve flag. We passed this output to the pedigree-based recombination map workflow as described below.

#### *Construction of a pedigree-based recombination map*

We used Lep-MAP3 (Rastas 2017) and pedigree data to construct a recombination map for the *Geospiza* samples on Daphne. We first filtered the GLIMPSE imputed dataset to only include variants with reference allele frequency (RAF) > 5% and sample allele frequency (AF) > 5%, as well as samples with > 0.0625 phasing accuracy (INFO field). We next used bcftools +prune to retain only 1 SNP every 1000bp and inferred parental genotypes using the Lep-MAP3 ParentCall2 command. As input, we passed 818 parent-offspring pairs to this command. We next assigned markers to linkage groups using SeparateChromosomes2 and joined singles to these linkage groups using JoinSingles2All, both with a lodLimit of 10. Finally, we ordered markers using OrderMarkers2 and the chromosome=1 flag. Positions were matched to the coordinates of Camarhynchus_parvulus_V1.1 using a custom awk script. The total genetic length in males was

2223cM and 2244cM in females.
In order to produce a linkage map to pass to SHAPEIT4 and GLIMPSE, we applied custom filtering. First, chr25 was split into two linkage groups by Lep-MAP3 and we combined these into one manually. This may have arisen either as the result of an assembly error. We next used GAM smoothing from the R package `mgcv` (Wood 2011) to build a smoothed recombination file across all positions (i.e. genetic position ~ physical position). We used the average genetic position of males and females for this analysis. For chromosome Z, we only include positions from the male map. We subsequently removed variant positions with residuals > 8 or < -8 derived from the GAM model.

The genmap format used by SHAPEIT4 requires that the genetic positions always vary positively with physical position. To force our linkage map to conform to this, we reversed the order of 18 chromosomal regions where the orientation of the Lep-MAP3 linkage map is reversed to genomic coordinates. These files were written to gmap format (i.e. one file per chromosome with physical and genetic coordinates) and are available as supplemental data. We additionally used the MareyMap R package (Rezvoy et al. 2007) to generate recombination rate estimates using a sliding window approach with a window size of 1Mb, a shift of 500kb, and threshold of 8.

***Reference panel and imputation with recombination map***

For our second phase of imputation, we reconstructed the reference panel using the newly generated recombination map. At this point, we generated sequencing data for an additional 92 individuals (see Table S1 and sampling above). We called variable sites using the same protocol as described above for the initial reference panel. We included all samples in a second run of SHAPEIT4, but this time by supplying the recombination map in .gmap format. Imputation in GLIMPSE was rerun using the same parameters as before but now including the recombination map in the GLIMPSE_phase step.

We evaluated imputation accuracy by selecting a high-coverage *Geospiza fortis* (approximately 20x coverage) and downsampling the mapped reads using samtools -s 1.02 to achieve an approximate coverage of 2x. We removed this individual from the reference panel and re-ran our GLIMPSE pipeline to impute genotypes. Concordance was evaluated for allele frequency bins of 0.00000, 0.00100, 0.00200, 0.00500, 0.01000, 0.05000, 0.10000, 0.20000, 0.50000 using GLIMPSE_concordance and visualized using the helper script concordance_plot.py.

We removed sites with sample minor allele frequency (i.e., among Daphne individuals) or reference minor allele frequency of less than 0.05. We also removed sites of imputation quality < 0.065 (i.e., the info field generated by GLIMPSE). The resulting filtered variant file with phased and imputed genotype calls was used for all downstream analyses of variant data. This dataset consists of 5,109,160 SNPs.

***Genetic sex determination***

We used sequencing depth to assign genetic sex to each individual. We extracted sequencing depth from chromosomes 4 and Z using the command samtools coverage (v1.10) (Li et al. 2009)). We used the ratio of the number of reads mapped to chromosome 4 to Z to assign sex because females are the heterogametic sex in birds. We identified genetically male individuals with a ratio of <1.2 chr4 to chrZ reads, and individuals with a ratio of >1.7 chr4 to chrZ reads were assigned female. In both cases, we required an average sequencing depth of 0.15. We could not assign 30 individuals using this cutoff (0.7% of samples). However, 16 of these individuals had been assigned a sex in the field based on breeding or morphology. The genetic sex of 164 samples (7.3% of samples) conflicted with the assignment in the field. Males and females cannot be distinguished in the field in immature (brown) plumage. In subsequent molts males, but not females, acquire increasing amounts of blackness in the plumage. Furthermore, only males sing. We previously confirmed the accuracy of sex identification using sex-linked markers (Enbody et al. 2021). The assigned genetic sex was used for all downstream analyses that include sex.

***Admixture and ancestry estimates***

We used ADMIXTURE v1.3.0 (Alexander et al. 2009) to estimate sample genomic ancestry from a set of ancestry informative and putatively neutral, unlinked markers. We generated ancestry-informative markers by calculating a four-population Population Branch Statistic (PBS) based on the formula in (Zhan et al. 2014). We calculated per site pairwise population divergence Fst using the software pixy (Korunes and Samuk 2021) for all combinations of *G. fortis, G. scandens*, *G. fuliginosa*, and *G. magnirostris* in the reference panel. We excluded samples of these species from Daphne (*n* = 26). We calculated the four-population PBS statistic using a custom script in R. A brief R chunk example is shown here for *G. magnirostris*:

```
PBS_magnirostris = (-log10(1-fst_fuliginosa_magnirostris) +
-log10(1-fst_fortis_magnirostris) +
-log10(1-fst_magnirostris_scandens) -
          -log10(1-fst_fortis_fuliginosa) -
-log10(1-fst_fuliginosa_scandens)) / 3)
```

We next identified sites exceeding the 95% confidence interval for each of the four species, *i.e.* loci at high frequency within the focal species that are low frequencies in others. We took this approach due to the high frequency of allele sharing among species of Darwin's finches (Lamichhaney et al. 2015). For this reason, we consider the markers to be "ancestry informative markers" and are not an unbiased and random selection of markers across the genome. We removed any position overlapping with the 28 loci reported in Rubin et al. (2022). The initial set of SNPs included 574,812 positions. Finally, we pruned variants within 20kb windows with an $R^2$ exceeding 0.3 using bcftools +prune, resulting in 156,977 unlinked and ancestry informative sites.

We performed a series of admixture runs from K = 2 to K= 10. For a given K, we first performed a training admixture run using an equal number (n = 56) of *G. fortis*, *G. fuliginosa*, *G. scandens*, and *G. magnirostris* samples to avoid biases associated with unequal sampling of species within our dataset. We then projected our full sample dataset onto the population structure (allele frequencies) inferred during the training run by invoking option –P (Shringarpure et al. 2016). For each value of K, we conducted 100 independent runs and summarized runs using CLUMPAK (Kopelman et al. 2015). We determined the optimum K based on cross-validation error.

***Heritability estimates***

We estimated SNP heritability using the GREML functions in the software GCTA (Yang et al. 2011). We only estimated SNP heritabilities for *G. fortis,* because large sample sizes are required for accurate SNP heritability estimates (Yang et al. 2014). Specifically, we estimated heritabilities for the relatedness grouping that includes *G. fortis* described in the methods below. To prepare the inputs for GCTA, we converted the VCF to binary ped format using qctool (v2.0.6, https://www.well.ox.ac.uk/~gav/qctool/) and used custom `sed` functions to rename chromosomes to numeric-only names temporarily. We generated a single input for each chromosome.

We built a linkage-disequilibrium (LD) and minor allele frequency (MAF) stratified relatedness matrix (LDMS) following the protocol described in (Yang et al. 2015). We stratified SNPs by estimating LD in regions of 200bp using the `--ld-score-region` flag in GCTA. The flag `--autosome-num` was required for this and all subsequent commands in GCTA due to the additional chromosomes in birds compared to humans. The Z-chromosome was excluded for all relatedness matrices generated here and the W-chromosome is not included in the assembly. We next stratified all SNPs into bins of high or low LD based on whether each SNP lay above or below the median LDscore_SNP value, respectively. These SNP sets were further stratified into two bins of high (maf > 0.05) or low (maf > 0.05) allele frequency. Four genomic relatedness matrices (GRMs) were constructed based on these SNP bins:

High-LD, High-MAF
Low-LD, High-MAF
High-LD, Low-MAF
Low-LD, Low-MAF

For each relatedness matrix, we passed the `--make-grm-alg 1` flag due to the inclusion of many closely related individuals in the dataset. The four relatedness matrices were then passed into a single run of GREML to estimate SNP-heritabilities of each SNP bin while taking into account the remaining three bins. SNP heritabilities were estimated in this way for beak size, beak shape, and weight for *G. fortis*. We inverse-normalized measurements in grams for body weight to improve normality. Including two principal components as additional covariates did not qualitatively change the results of these estimates.

To explore the effect of relatedness on the estimates from GREML we additionally built GRMs for each chromosome separately. For this analysis, we did not stratify SNPs into bins and instead included all variants in the dataset. We ran GREML for each chromosome separately and reported the sum of V(g)/Vp across all chromosomes.

We also estimated SNP heritabilities after identifying the six QTLs in the Genome-wide association analysis section below. For this analysis, we built a GRM for all autosomal SNPs, excluded SNPs with a MAF < 0.05, and included the `--make-grm-alg 1` flag. For each of the three phenotypes (beak size, beak shape, and body weight) we ran GREML with a set of covariates that included the genotypes of each of the six large effect loci. We estimated $V_{QTL}$ as the difference in V(g) estimated by GREML between estimates derived from GREML without any covariates compared to the one with covariates. i.e. $V_{QTL} = V(g)_{base} - V(g)_{QTL}$.

One method for reducing the impact of closely related individuals on SNP-heritability estimates for datasets containing many closely related individuals is to remove the closest relatives from the dataset before running the analysis. The number of off-diagonal individuals with a relatedness coefficient greater than 0.05 (commonly applied to human samples) removed all but 31 individuals. Due to this high relatedness, we could not partition our dataset into unrelated and related to assess the difference in $h^2_{SNP}$ and $h^2_{Pedigree}$.

***Genome-wide association analysis***

We conducted genome-wide association analysis (GWAS) using the software GEMMA (v0.98.4, (Zhou and Stephens 2014). Species of finches on Daphne are known to hybridize and their field species designation (described above) may not necessarily capture genetic relatedness. For this reason, we initially separated all individuals on Daphne into genetic clusters to reduce the potential confounding effects of cryptic ancestry on GWAS inference. A relatedness matrix of each ancestry cluster was used as a covariate for all GWAS run in GEMMA.

We constructed a GRM using all Daphne individuals of the four ground-finch species included in this study (*n* = 3,958). We generated a set of LD-thinned SNPs by selecting one SNP every 20kb using bcftools +prune (v1.10)(Danecek et al. 2021). We converted this VCF to bimbam format using the software qctool. We built a relatedness matrix using GEMMA with the `-gk 1` method. We supplied this relatedness matrix to the `hclust` function of R (v4.1.1, ) and `cutree` with *k* = 3 groupings to group samples into the three main ancestry groups. We selected *k* = 3 for this analysis after initial exploration (and the admixture analysis above) confirmed the very close genomic relationship among *G. fortis* and *G. fuliginosa* samples. All *G. fuliginosa* samples group within the *G. fortis* cluster using this method were removed from the GWAS due to their small sample size. We also excluded samples 01Dap21274 and 01Dap21269 as we suspected immigrants from Santa Cruz Island. The three relatedness clusters were subsequently labeled *G. fortis*, *G. scandens*, and *G. magnirostris* based on the majority membership of each species of each cluster. A separate relatedness matrix (*n* = 3) was generated for each species cluster and used for the downstream analysis of each species separately.

For each run of GEMMA we included the relatedness matrix, a phenotype file and a list of covariates. We focus on three key phenotypes: beak size (PC1), beak shape (PC2), and body weight. Morphological PCA was conducted separately for each relatedness cluster described above. To compare effect size directions among species, we forced a positive correlation among species for the eigenvectors representing beak size and beak shape. Specifically, we ensured a positive correlation between beak size and the sum of all three beak dimensions and between PC2 and the ratio of beak length to beak depth. In other words, we enforce that each PC is positively correlated with beak morphology across species. For body weight, we inverse-normalize transformed raw measurement values. GWAS were conducted in GEMMA for all three species *G. fortis*, *G. scandens,* and *G. magnirostris* under the following conditions:

1) A multivariate analysis using both beak size and beak shape as response variables. Both sex and body weight were included as covariates, in addition to the GRM for the focal species.

2) A univariate analysis using body weight as the response variable and sex as a covariate, in addition to the GRM for the focal species.

For *G. fortis,* additional analyses were conducted in GEMMA:

1) A multivariate analysis using both beak size and beak shape as response variables. In this analysis, only sex (but not body weight) was included as a covariate, and the GRM for the focal species.
2) A univariate analysis for beak PC1 using sex, body weight, and the highest -$\log_{10}$ *P*-value SNP on chromosome 1A from the multivariate analysis. This analysis evaluated the extent of linkage disequilibrium in the large QTL on chromosome 1A around the *G03* locus.

The first analysis was run to investigate the effect of including body weight as a covariate in the analysis. Five beak morphology loci are not significant in the body weight GWAS, indicating that they are unlikely the result of collider bias, *i.e.,* false associations for beak size by including body size as a covariate (Aschard et al. 2015) (Fig. S9).

For *G. fortis*, we also conducted a leave-one-out-chromosome (LOCO) analysis in GCTA for beak size, shape, and body weight. As mentioned above, for beak traits, we included body weight as a covariate and sex was included as a covariate in all runs. In a LOCO analysis, we built the relatedness matrix by excluding the focal chromosome to minimize the impact of QTLs on the focal chromosome on calculating associations. In general, we recovered the same six loci identified in GEMMA. The main difference is much broader QTLs and additional broad signals on chromosomes 3 and 5 for beak shape and beak size, respectively (see Supplemental Text 4).

To establish a cutoff value for significance, we permuted phenotypes using beak size (PC1) using the tools built into gemma-wrapper. We ran for 50 iterations to establish a strict threshold of -$\log_{10}$ *P*-value = 7.7 (and lax threshold of -$\log_{10}$ *P*-value = 6.6) for testing the statistical significance of GWAS hits. This value is approximately consistent with a Bonferroni corrected *P*-value of 0.05.

#### ***Haplotype classification at GWAS loci***

To identify the haplotypes associated with each QTL signal, we clustered individuals into groupings based on the top associated SNPs identified in our GWAS analysis. We first selected all variants surpassing the threshold of -$\log_{10}$ *P*-value = 6.6 and defined QTLs as genomic intervals containing outlier SNPs within 75kb of another outlier SNP. We created interval ranges using GenomicRanges (Lawrence et al. 2013) in R (v4.1.1). We further pruned these intervals to retain only intervals with SNPs, surpassing the strict threshold set by permutation of -$\log_{10}$ *P*-value = 7.7 and removed intervals spanning a length <100kb. This summary was performed for each GEMMA GWAS described in this study.

For the main regions of associations in the *G. fortis* multivariate GWAS (*i.e.*, using beak size and beak shape) we extracted the top 100 variants from each locus defined above. We clustered samples using an interactive process, beginning with *G. fortis* and expanding to other species. For each locus, we first use all 100 outlier genotypes among the *G. fortis* samples (within the genetic grouping defined above) and conduct a PCA ($n$ = 1,508) using 'prcomp' in R (v4.1.1). Samples were clustered into three groups using 'kmeans' in R based on genotypic PC1 by picking 25 samples at random and iterating 100 times. We next calculated allele frequencies among the three groups and selected the top 5 SNPs most strongly differentiated among the groups with the lowest and highest PC values (*i.e.*, alternative homozygotes). We used these five SNPs to rerun PCA and kmeans clustering on the full dataset to identify haplotypes in all finches analyzed here. We initially identified the heterozygous grouping as the intermediate grouping in the genotypic PC1 and confirmed this using haplotype plots for the main six loci considered in this study. At this point, we calculated (among *G. fortis*) the average bill size in each of the three groupings and applied the label "S" to the smaller haplotype and "L" to the largest haplotype (individual genotypes are "LL", "LS", or "SS").

We applied custom curation to four of the loci. For locus *G01, G03, G07,* and *G30* we noticed a number of *G. scandens*-specific variants that separated a third haplotype. At *G03*, this second locus most closely resembled the haplotype most common in *G. magnirostris*. Correspondingly, for *G03* we identified variants within the locus interval with high allele frequency variation among *G. scandens* and *G. magnirostris* and used these variants to separate a second L haplotype (L1 and L2). This haplotype variation corresponds to the two L haplotypes described in (Lamichhaney et al. 2018). For *G07*, the *G. scandens* haplotype most closely resembled the *G. fuliginosa* (pointed) haplotype so we used high delta allele frequency variation SNPs between these two species in the *G07* interval to bin individuals for a third pointed haplotype (P1 and P2). For these two loci the third haplotype was binned with L and P, respectively, for all effect size estimates in the main text.

For *G01* and *G30* we used all 100 top SNPs to classify individuals as *SS*, *SL*, and *LL*, but noted some heterogeneity in variants specific to *G. scandens.*

#### ***Effect size per locus***

We estimated locus-specific effect sizes using the haplotypes described above. Effect sizes were estimated separately for the genetic groupings of *G. fortis, G. scandens,* and *G. magnirostris.* For each locus, we conducted a type III analysis of variance (ANOVA) using the R package 'car' (Fox and Weisberg 2011) using bill size (PC1), bill shape (PC2), or body weight as the response variable. We used all six loci as the predictor variables and computed the partial effect size eta-squared ($\eta^2$). This calculation estimates the variance explained by each predictor variable after considering the variance explained by all other predictor variables. We also re-calculated effect sizes for the residuals of a model with bill size or bill shape as the response variable and weight and sex as the predictor variable. This estimates the effect of each locus on beak size and shape independent of body size and sex.

#### *Genomic PCA*

We used plink2 to calculate a genome-wide PCA for all Daphne individuals. We compared genomic PC1 and genomic PC2 from this calculation to a PCA estimated from PCangsd v.0.982 (Meisner and Albrechtsen 2018) that used genotype likelihoods as input. Genotype likelihoods were estimated using angsd v.0.3 (Korneliussen et al. 2014) using the following commands:

```
$ANGSD -b $BAM_LIST \
      -uniqueOnly 1 -remove_bads 1 -only_proper_pairs 0 -trim 0 \
      -GL 2 -doMajorMinor 1 -doGlf 2 -doMAF 1 -minMapQ 20 -minQ 20 \
      -SNP_pval 1e-6 -minMaf 0.05 \
      -out $OUT_PREFIX
```

Using a random 1% subset of SNPs extracted from the angsd derived genotype likelihood file, we then ran PCangsd as follows:

```
pcangsd.py -beagle $INPUT_BEAGLE -iter 1000000 -o $OUT_PREFIX
```

#### *Comparison to random loci*

We selected 100 random loci with starting allele frequencies equal to the frequency of each locus identified in the the GWAS in *G. fortis*. We accomplished this by first estimating the allele frequency of the six loci in the *G. fortis* and *G. scandens* populations in 1983. Next, we excluded SNPs inside these six association regions and the 28 regions identified in Rubin et a. (2022). From these remaining SNPs, we selected 100 random SNPs within 0.01 frequency of the starting frequency for each lous. Thus, for each species, we compiled a list of 100 random SNPs per locus with a starting allele frequency similar to the six loci identified in the study. This corresponds to 600 loci for *G. fortis* and 600 for *G. scandens*.

#### *Drought analysis*

We selected the 71 *G. fortis* samples alive before the 2004/2005 El Niño drought event for refined analysis of survival during this period. 37 individuals survived and 34 died. We tabulated the number of individuals at each locus and genotype class that survived. Next, we estimated selection coefficients as described (Hedrick 2011, 4th edition, p. 144).

To assess the contribution of each locus to survival, we summed for each individual the number of copies of the small allele they carried (*i.e.* SS = 2, SL = 1, LL = 0, where S is the small allele and L is the large allele). To determine if any locus does not contribute to the overall survival advantage, we tabulated this value from leaving one locus out at a time. On each of the resulting values of ($n$ = 5 + the full model), we included the number of small alleles as the predictor variable for a generalized linear model in *R* with survival as the response variable. We compared the Akaike information criterion (AIC) for all five models compared against the full model (*i.e.,* one that included all loci) and determined that removing GF07 and GF30 reduced the overall AIC score. These two loci were dropped, and the final model results were presented for the four remaining loci and plotted in Fig. 4.

### Supplemental figures

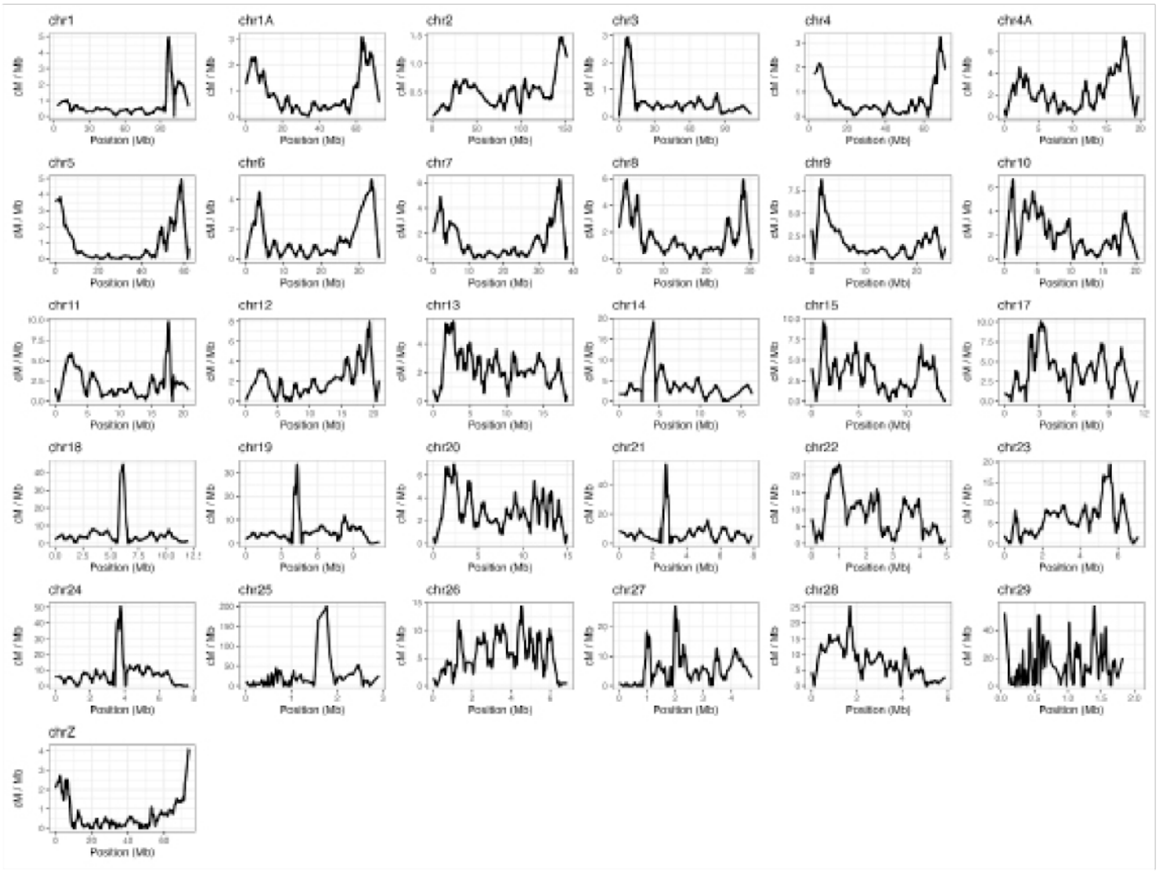


**fig. S1**: *De novo* pedigree-based recombination map based on *Geospiza* families on Daphne. The sex-averaged recombination rate is shown. The total genetic length in males was 2,223cM and 2,244cM in females.

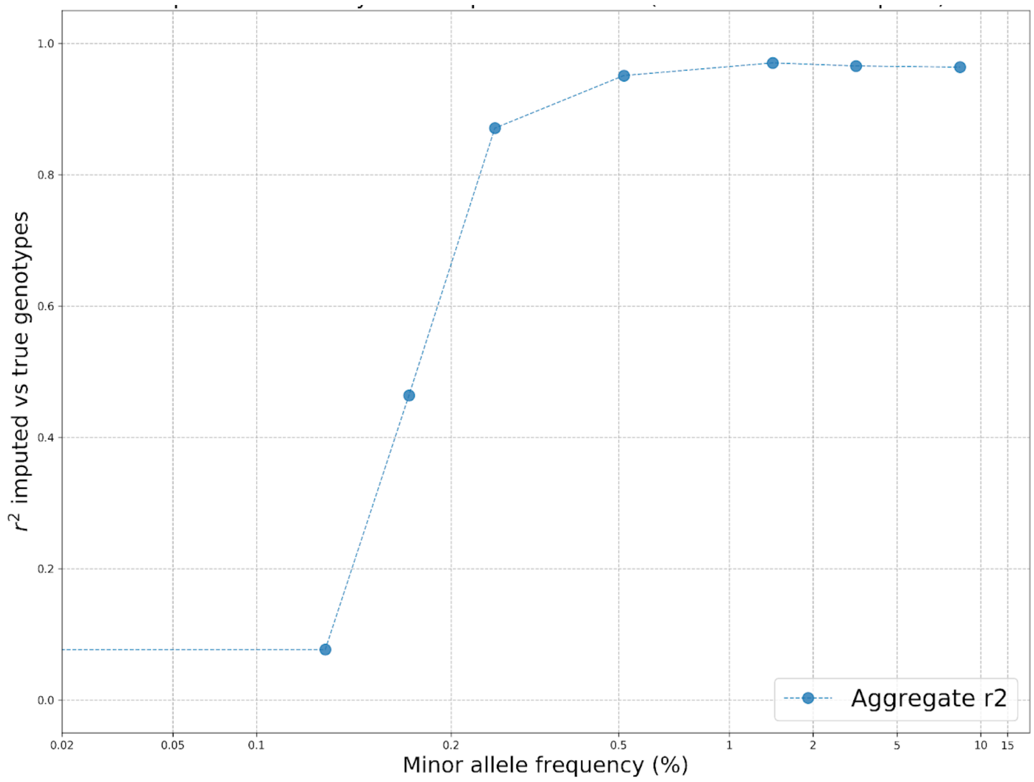


**fig. S2**: Imputation accuracy for a single down-sampled individual (5560) across the allele frequency spectrum. On the y-axis, the $r^2$ value for imputed true genotypes was calculated based on running imputation on a single individual after it was removed from the imputation panel.

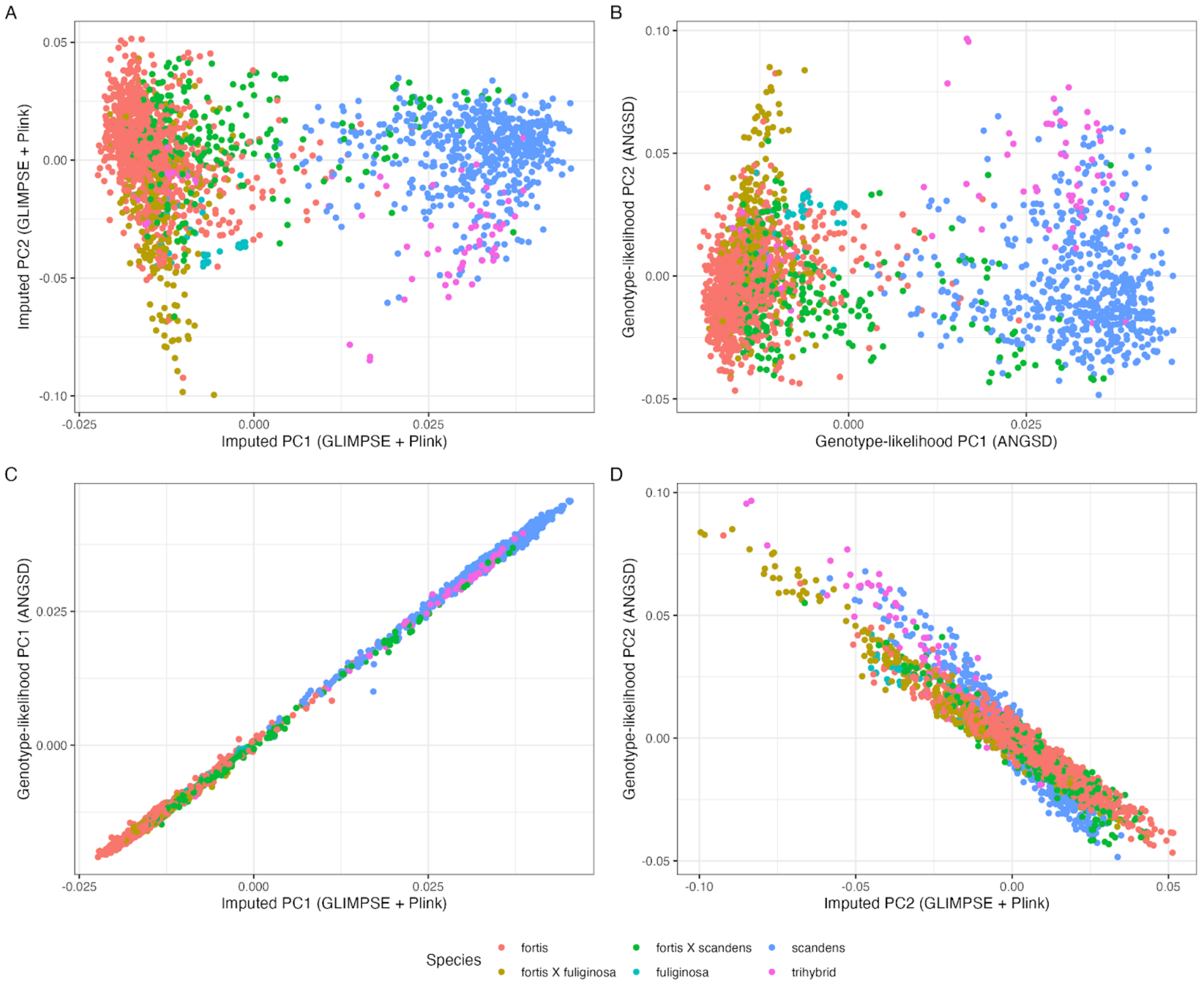


**fig. S3**: (A) Genomic PC1 and genomic PC2 from a PCA analysis using the imputed genotypes produced from the software GLIMPSE for all Daphne finches. (B) Genomic PC1 and genomic PC2 from a PCA analysis using a genotype likelihood framework in the software ANGSD and PCangsd. (C) Concordance between genomic PC1 from imputed genotypes (x-axis) and genomic PC1 calculated using PCangsd (y-axis). (D) Concordance between genomic PC2 from imputed genotypes (x-axis) and genomic PC2 calculated using PCangsd (y-axis). In all panels, colors refer to species assigned from field capture and pedigree analysis.

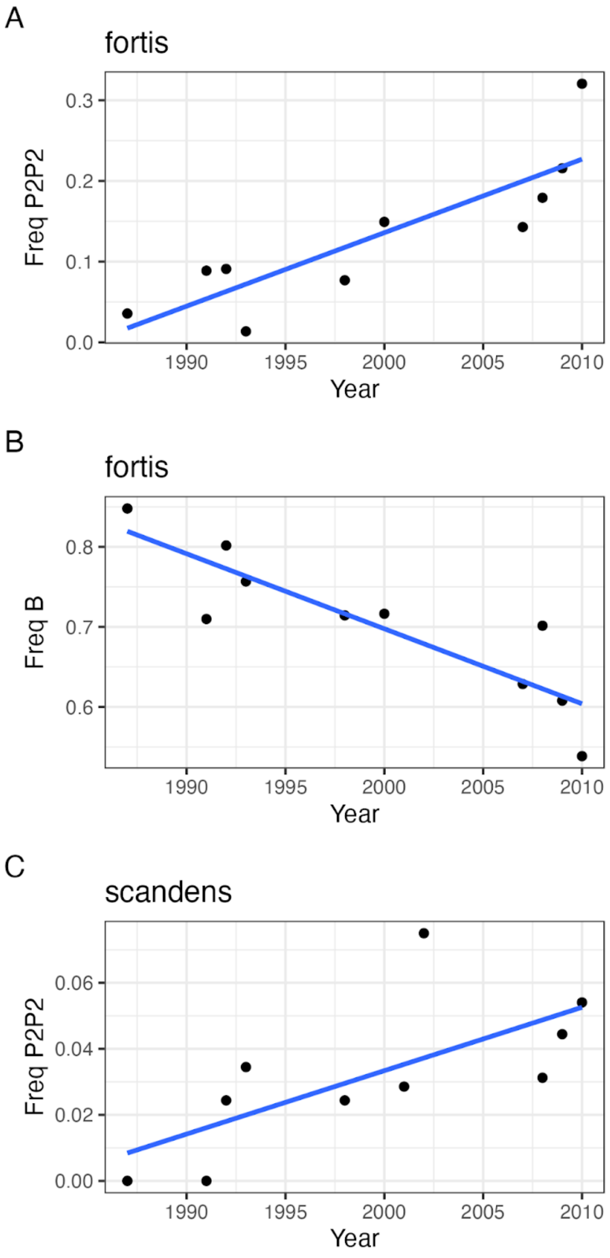


**Fig. S4**: Changes in haplotype frequencies in (a) G. fortis P2P2, (b) G. fortis combined haplotypes containing a B allele, and (c) G. scandens P2P2.

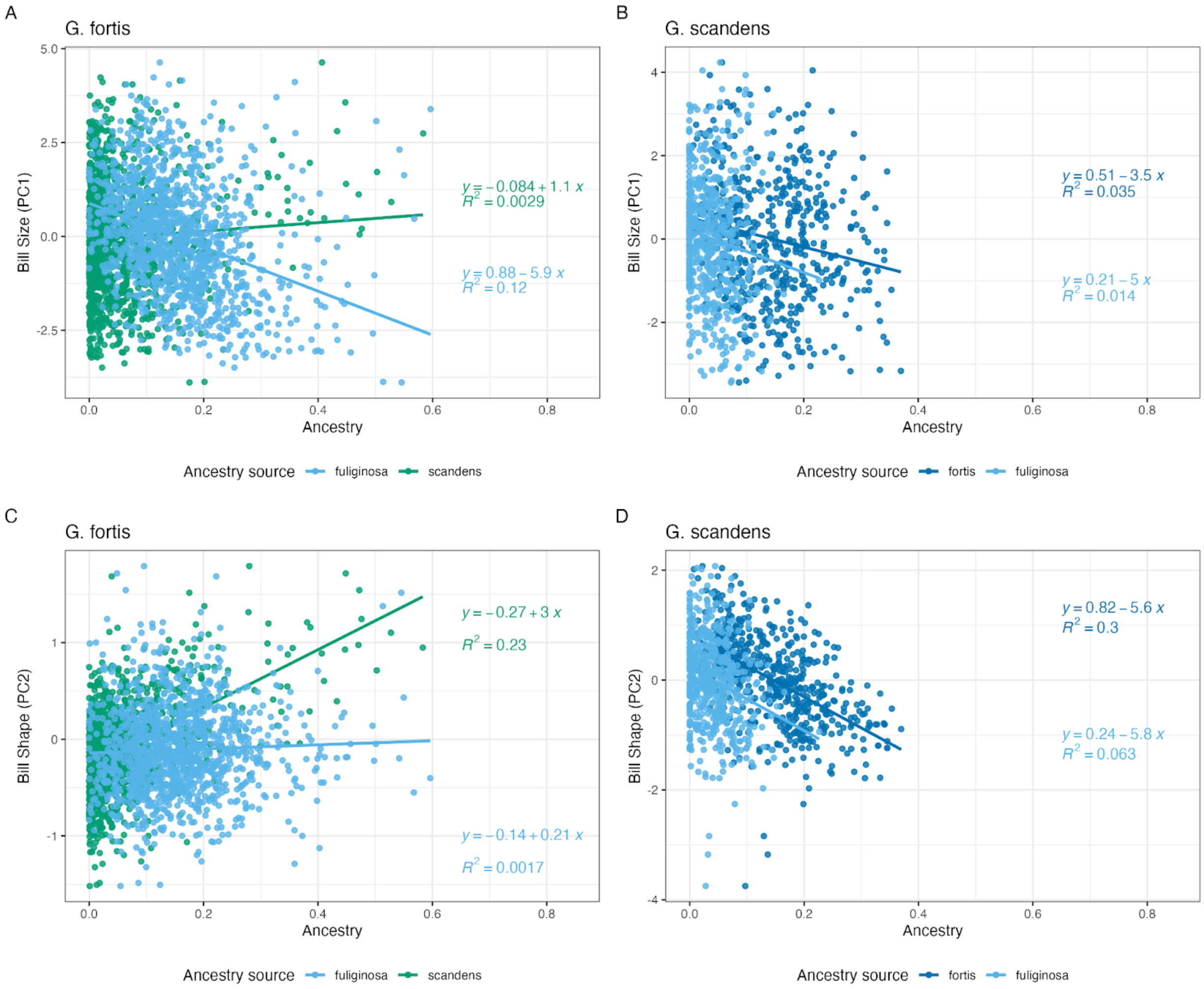


**Fig. S5**: A) *G. fortis*: The association between bill size and minor ancestry from *G. scandens* and *G. fuliginosa.* B) *G. scandens*: The association between bill size and minor ancestry from *G. fortis* and *G. fuliginosa.* C) *G. fortis*: The association between bill shape and minor ancestry from *G. scandens* and *G. fuliginosa.* D) *G. scandens*: The association between bill shape and minor ancestry from *G. fortis* and *G. fuliginosa.* Points and linear fits are colored by minor ancestry source.

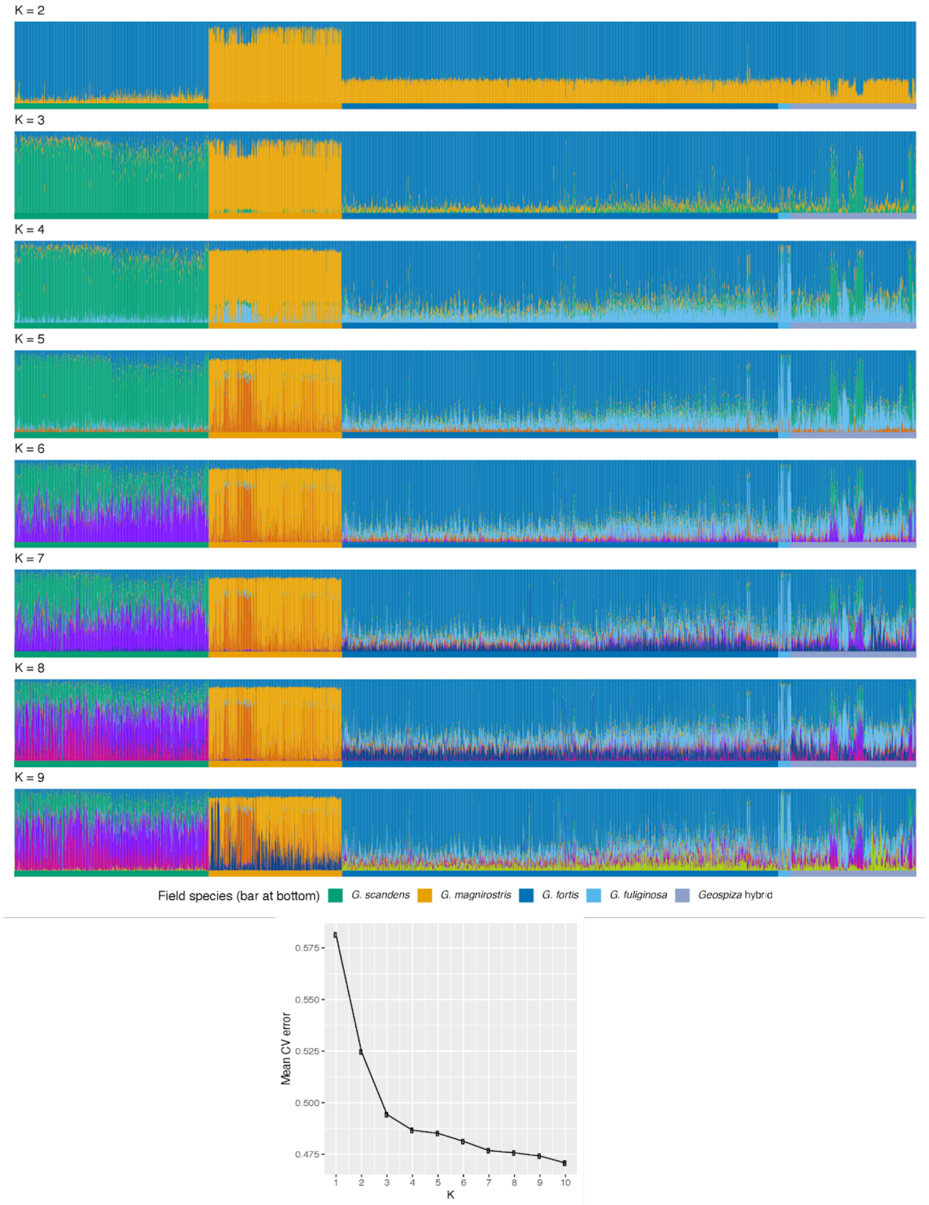


**fig. S6**: Admixture proportions for $K$ = 1-9 for all samples on Daphne. The ordering differs from Fig. 1 by grouping individuals by their field identification (bottom colored bar) rather than the relatedness matrix. A cross-validation curve for best fitting $K$ is shown for 1-10, with the best fitting value approximated at $K$ = 4 or $K$ = 5.

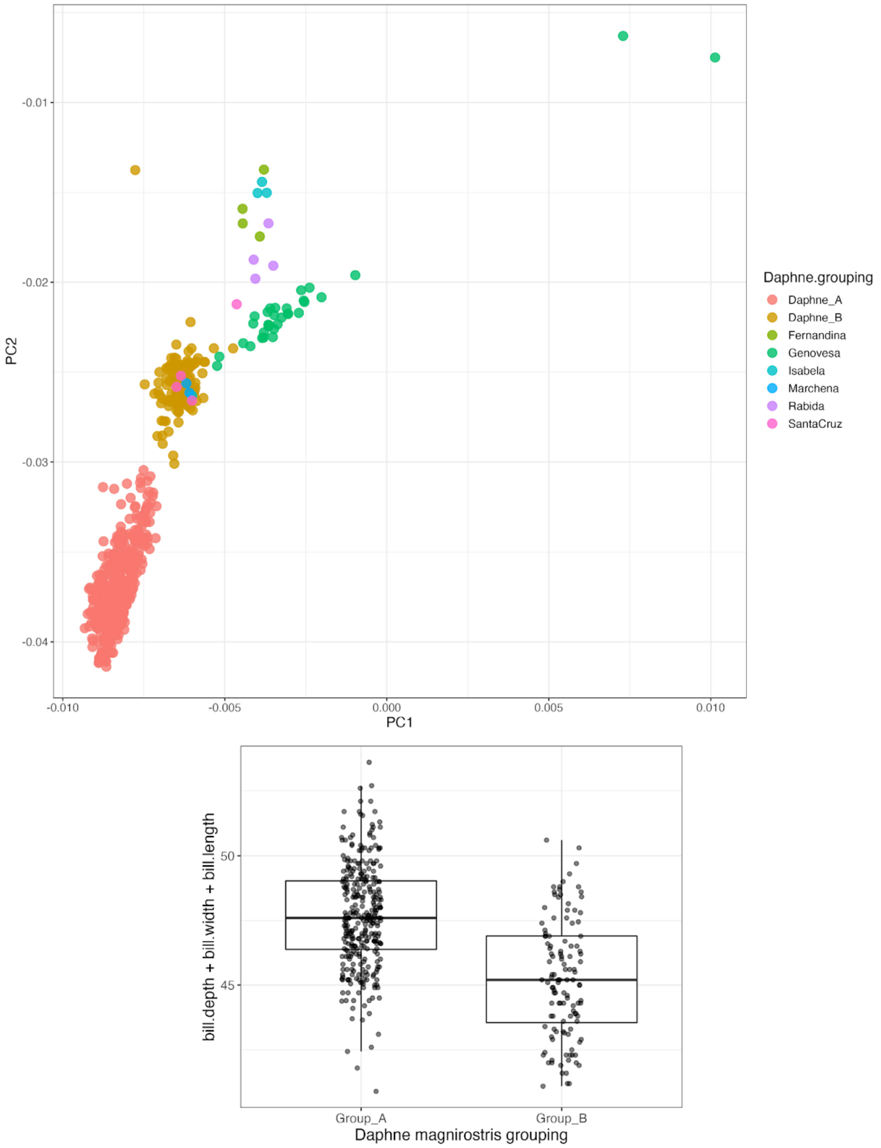


**fig. S7**: Top, Genomic PCA for all *G. magnirostris* samples in relation to all *G. magnirostris* present in the reference panel. Below, comparison of beak size (sum of all three dimensions) for the two *G. magnirostris* ancestry groups on Daphne. Group B is smaller and more similar to individuals from the nearby islands of Santa Cruz, Marchena, and Isabela. The island of origin for group A individuals is uncertain, but the most likely candidates are unsampled populations on Pinta and Santiago.

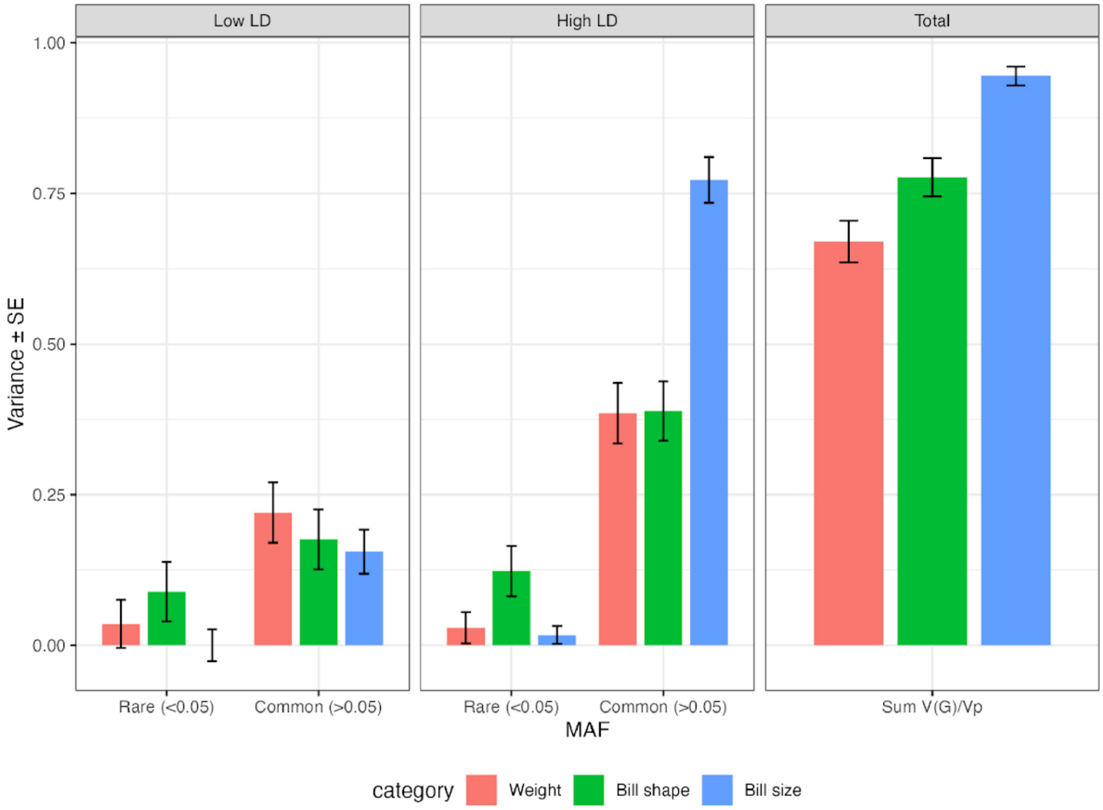


**fig. S8**: SNP heritability (V(G) / Vp) for three different phenotypic traits in *G. fortis*. Heritability is stratified by both allele frequency (<0.05 and > 0.05) and linkage disequilibrium. Linkage disequilibrium was stratified by greater or lesser than the mean SNP-LD score calculated using GCTA.

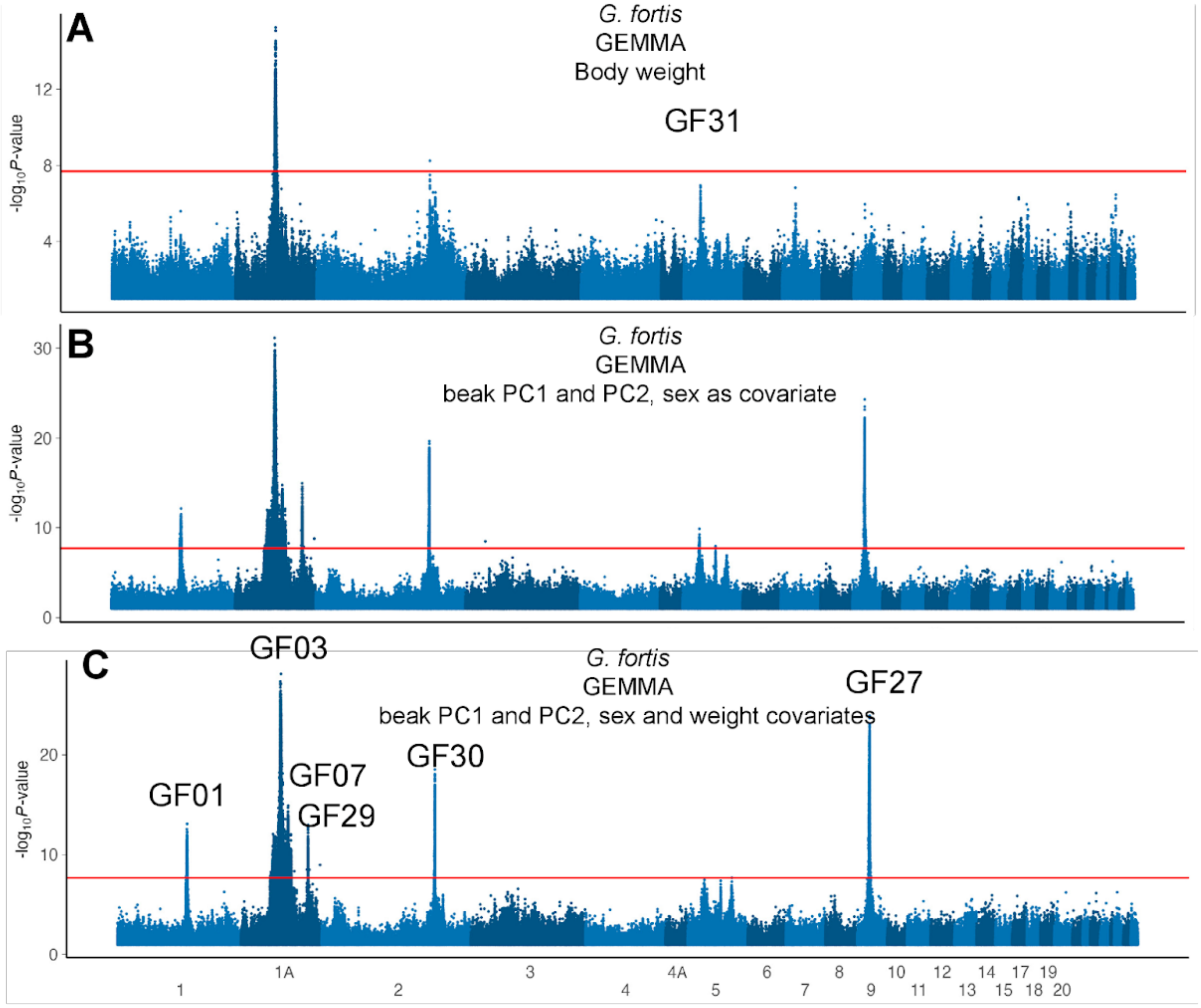


**fig. S9**: Genome-wide association analysis for three analyses performed in *G. fortis*. In (A), only body weight is included as a response variable, with sex as a covariate. In (B), beak traits P1 and PC2 are the response variables and sex is the only included covariate. In (C), beak traits P1 and PC2 are the response variables and both sex and body weight are included as covariates. The six loci shown in Fig. 2 are labeled, plus G31.

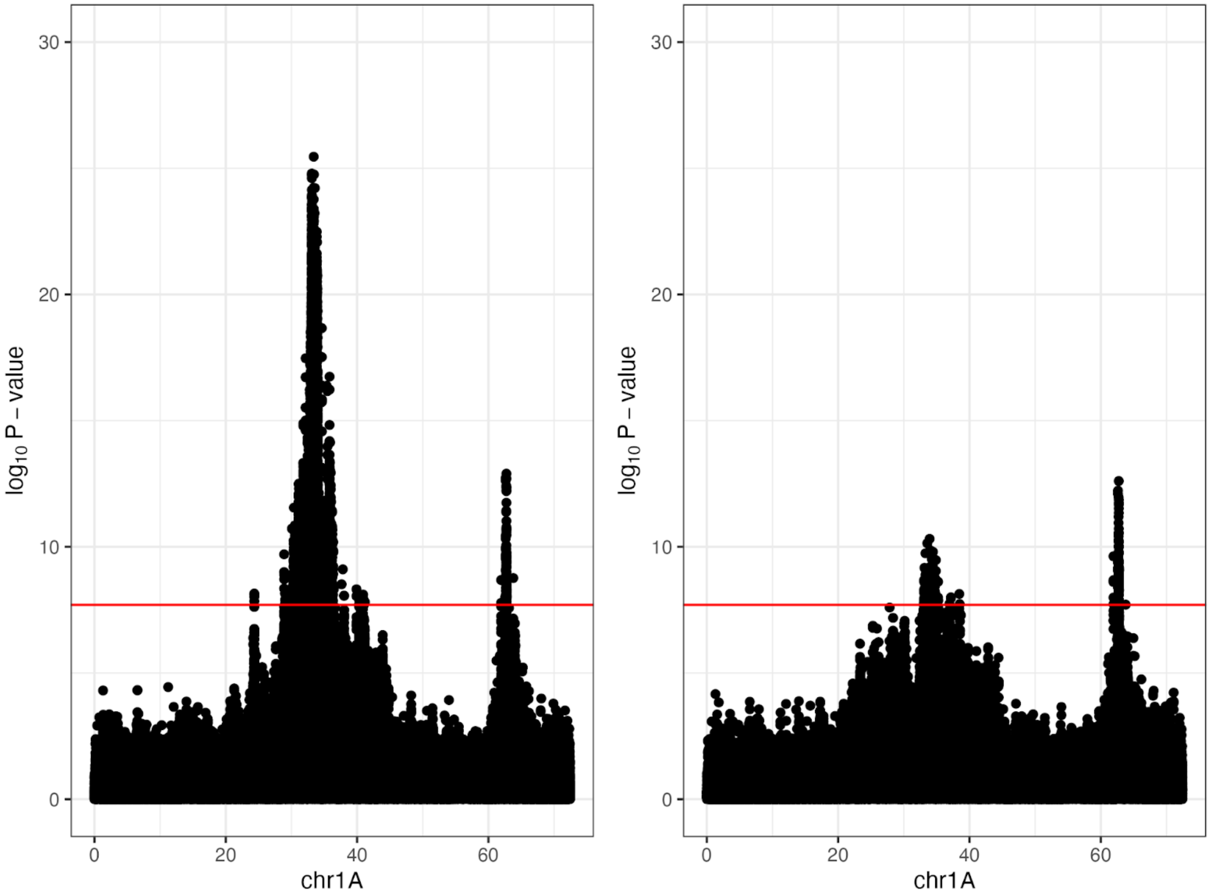


**fig. S10**: Left, a closeup of chromosome 1A from Fig. 2B. Right, GWAS using the top SNP in the locus *G03* as a covariate. The result demonstrates that most of the large region of association falls below the significance threshold used ($\log_{10} P = 7.7$).

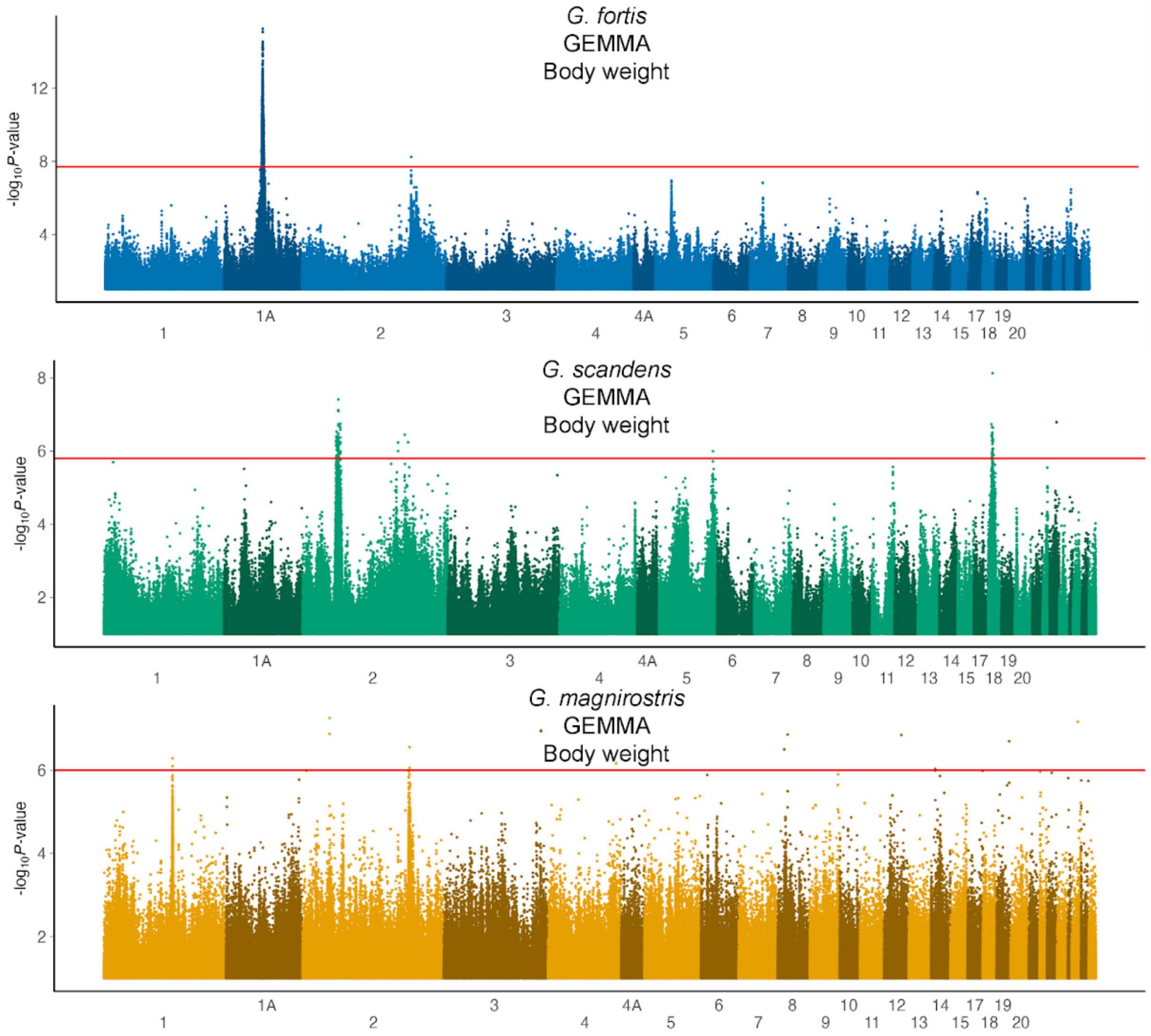


**fig. S11**: Genome-wide association analysis for body weight (sex as a covariate) for *G. fortis, G. scandens,* and *G. magnirostris.* A significance threshold of $\log_{10} P = 7.7$ is shown as a red line.

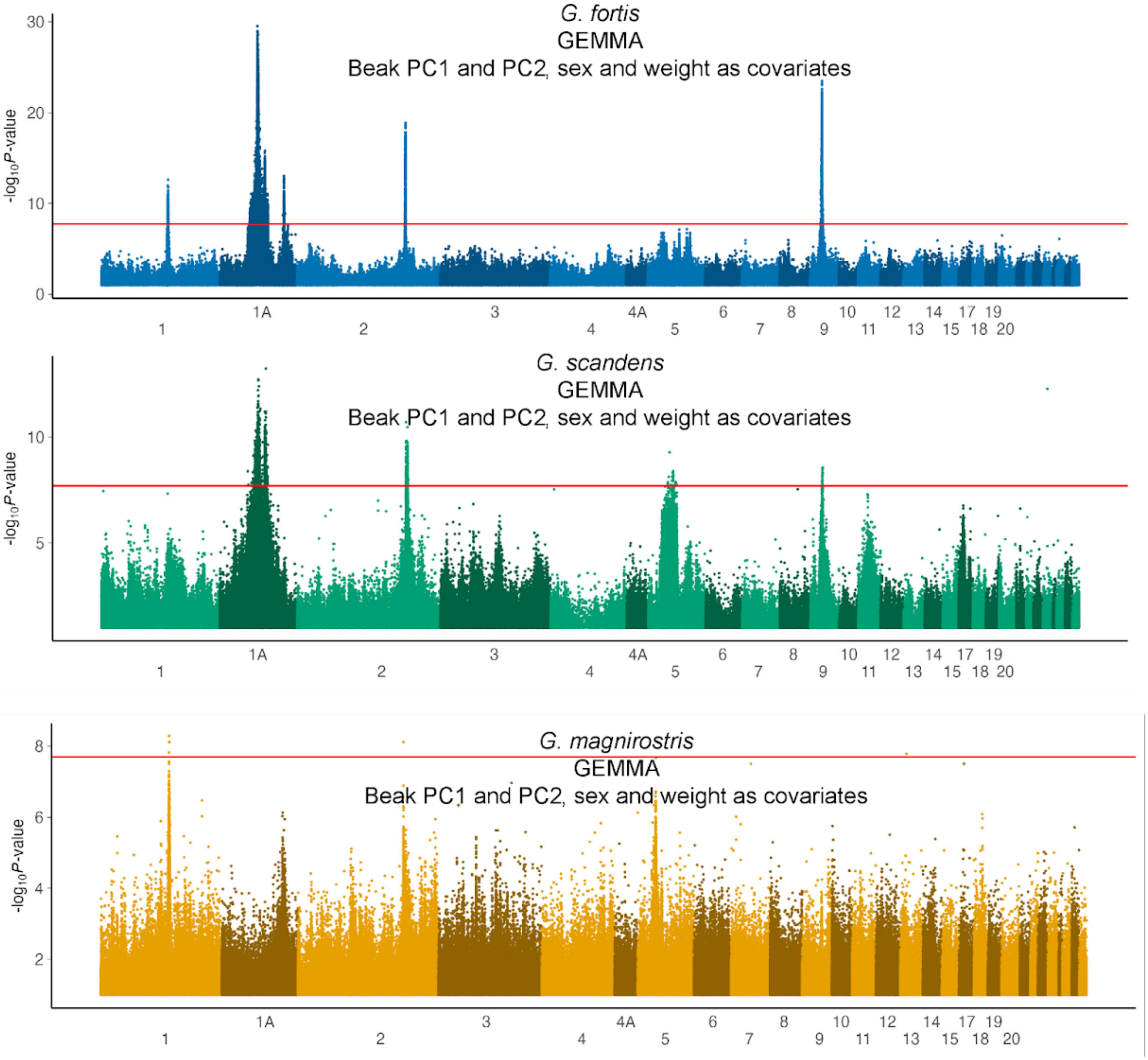


**fig. S12**: Genome-wide association analysis with both beak size (PC1) and beak shape (PC2) as response variables (sex and body as covariate) for *G. fortis* (same as Fig. 2)*, G. scandens,* and *G. magnirostris.* A significance threshold of $\log_{10} P = 7.7$ is shown as a red line.

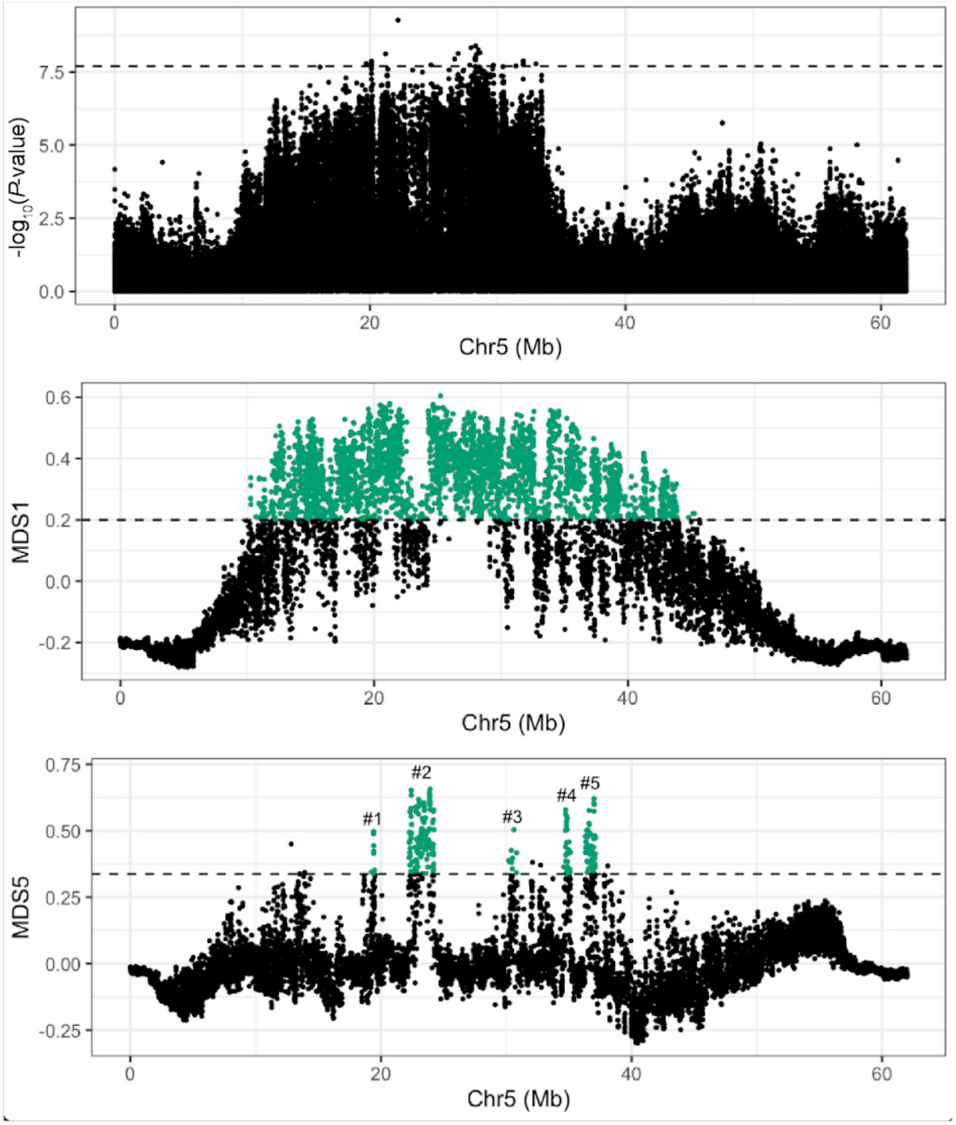


**Fig. S13**: **Top**, Association signal on chr5 for *G. scandens* shown in Fig. S12. **Middle**, multidimensional scaling (MDS) coordinate 1 generated in LocalPCA showing long-distance structuring of haplotypes in the region. **Bottom**, MDS coordinate 5 showing genomic regions likely representing recombination among haplotypes segregating in this region.

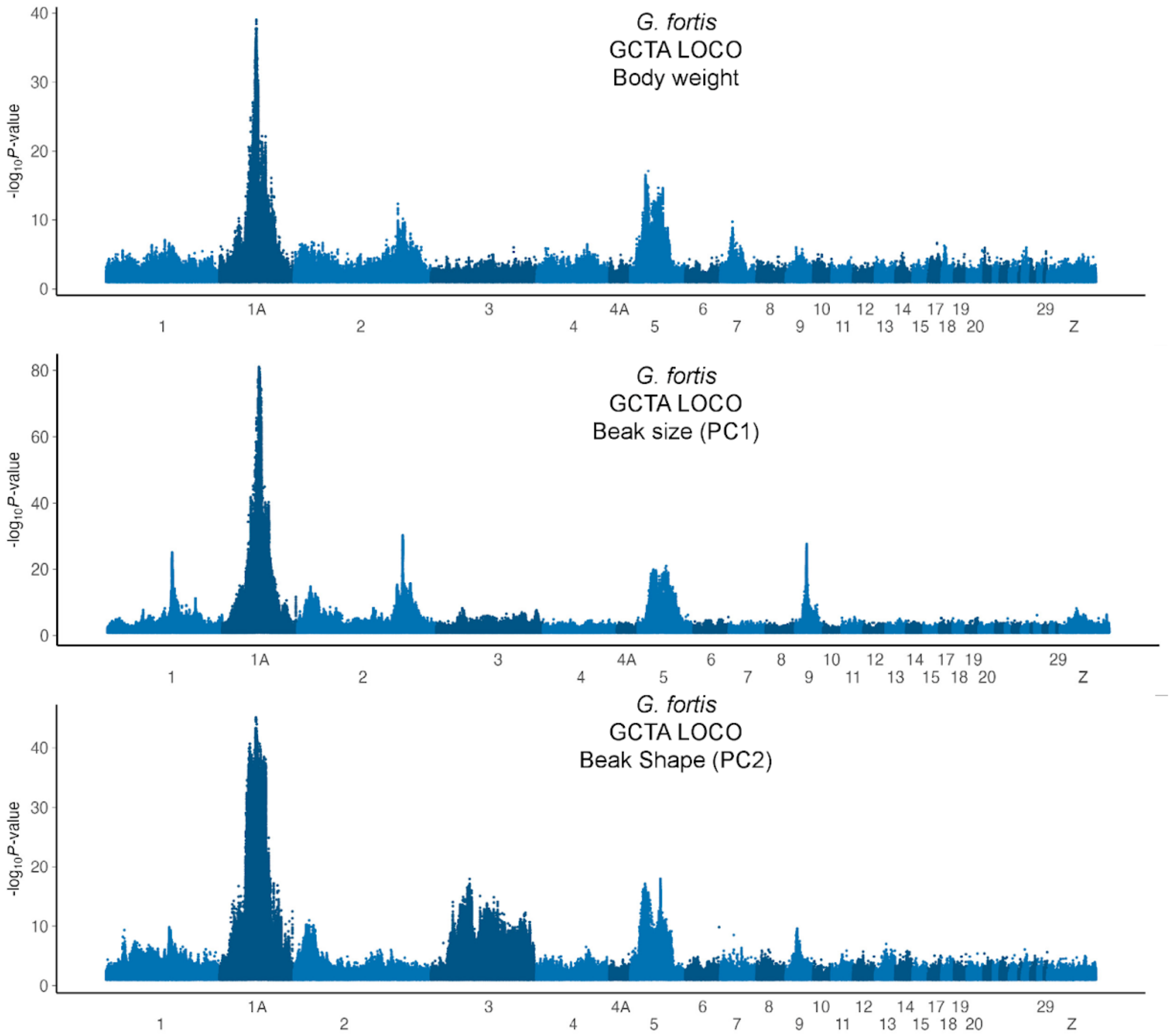


**fig. S14**: The results of a leave-one-out-chromosome analysis performed using the software GCTA for three phenotypic traits in *G. fortis*: body weight, beak size, and beak shape. The difference compared to other GEMMA analyses is that for each chromosome, a relatedness matrix is constructed by leaving the focal chromosome out. As a result, shared relatedness on that chromosome will improve the power for detecting QTLs. We interpret the broader peaks compared to GEMMA analyses shown elsewhere due to this analytical difference.

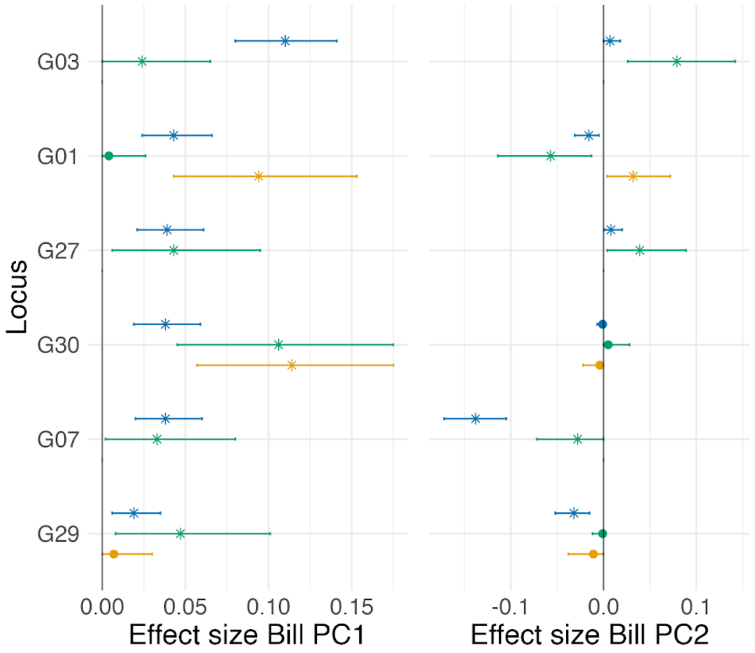


**fig. S15**:  Effect sizes for beak size and shape residuals. As in Fig 3, estimated effect sizes for all six loci shown in this study, plus the ten remaining loci identified in (Rubin et al. 2022). Here, effect sizes are estimated from the residuals of a model built using both sex and body weight as a covariate in the model. The variance explained for *G. fortis* is 29% for beak size and 20% for beak shape.

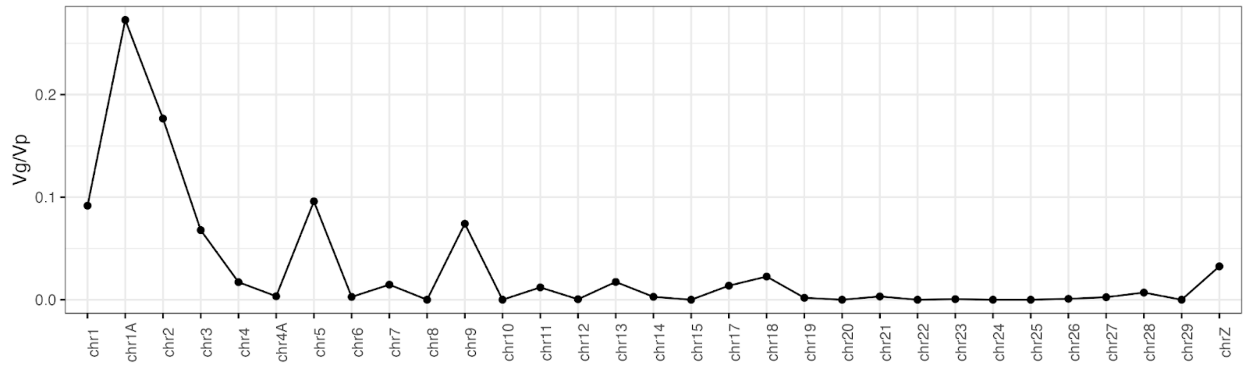


**fig. S16**: SNP Heritability (Vg/Vp) per chromosome where a relatedness matrix was built for each chromosome separately and each included as co-factors in a single GREML analysis for beak size in *G. fortis*. In other words, the values shown here are the SNP heritability score for each chromosome while considering estimates from all other chromosomes.

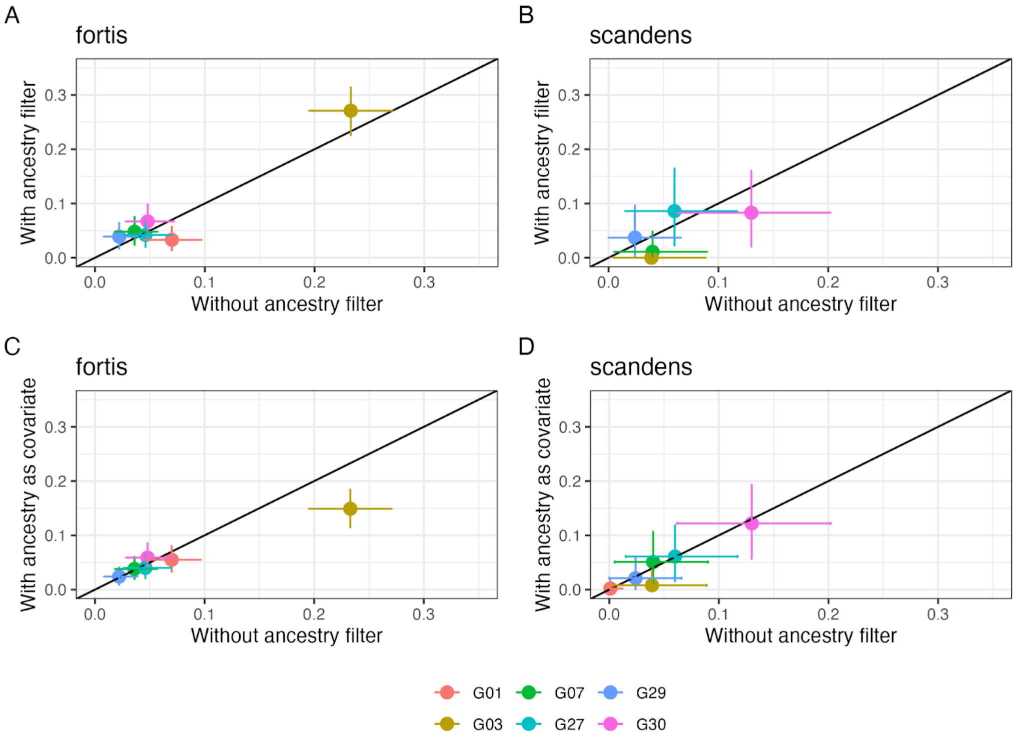


**fig. S17**: Effect size predictions for six loci identified in the *G. fortis* GWAS. (A-B) The y-axis shows effect size estimates described in the paper, and the x-axis shows effect sizes after removing individuals with self-ancestry values < the 75% quantile. C) *G. fortis* — y-axis: effect size estimate when including minor ancestry of *G. fuliginosa* and *G. scandens* as fixed effects. D) *G. scandens* — y-axis: effect size estimate when including minor ancestry of *G. fuliginosa* and *G. fortis* as fixed effects.

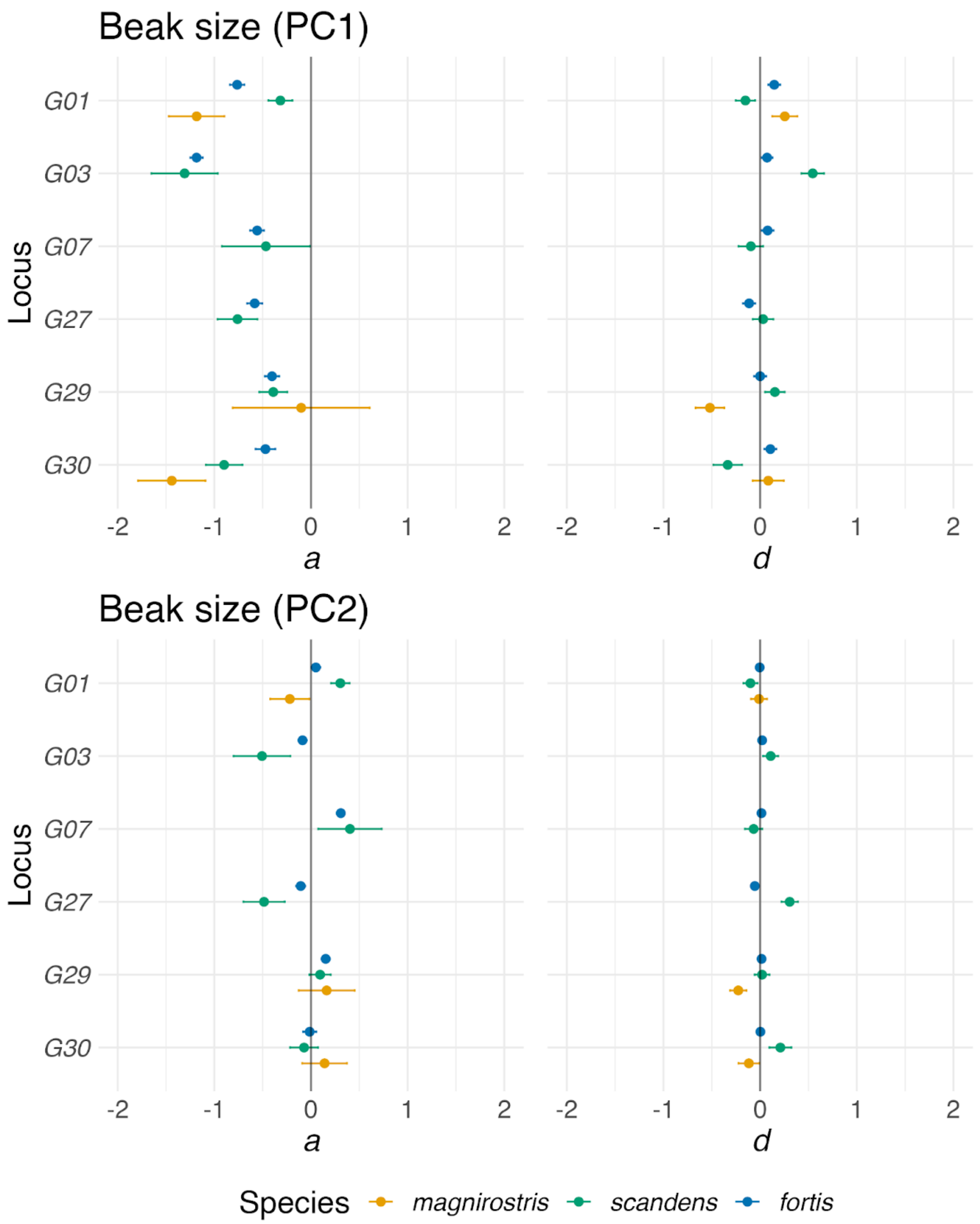


**fig. S18**: Additive (*a*) and dominance effects (*d*) of each of the six loci discussed in this study. For each locus, *a* and *d* are calculated relative to the small allele; i.e., negative values imply a smaller phenotype for homozygous small allele individuals (*a*) or the heterozygote is more similar to the small allele (*d*).

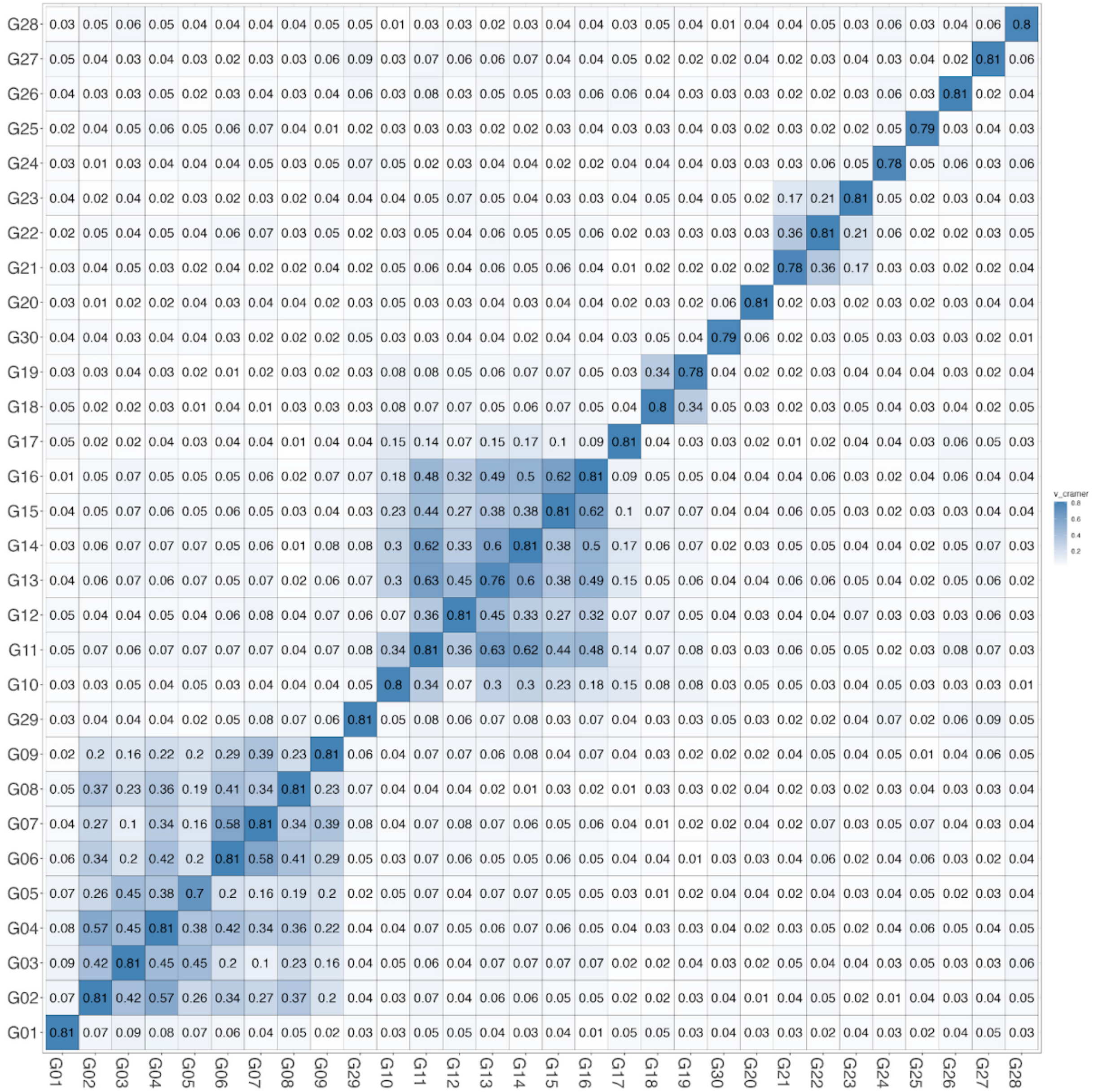


**fig. S19**: Linkage disequilibrium estimated using Cramer's V for all loci in this paper and those described in (Rubin et al. 2022). We considered locus pairs with a coefficient greater than 0.3 to be in disequilibrium.

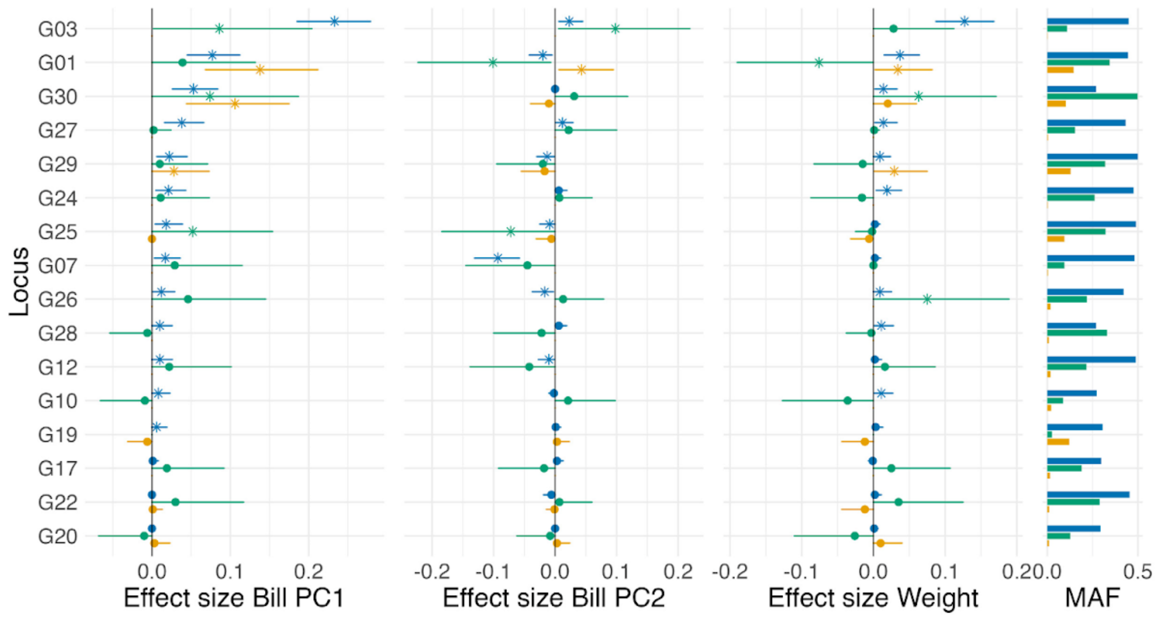


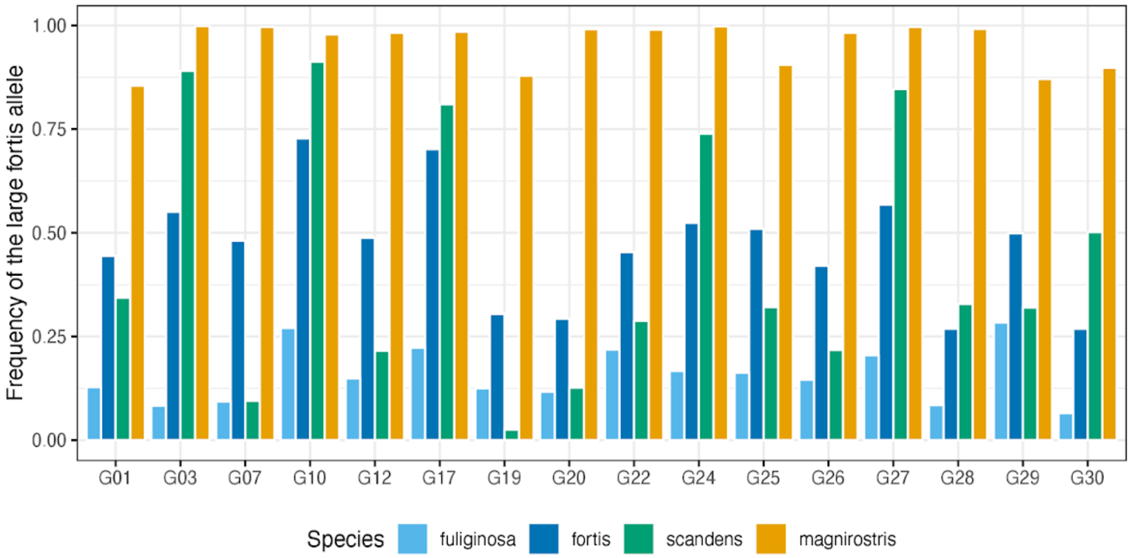


**fig. S20**: Top, as in Fig 3, estimated effect sizes for all six loci shown in this study, plus ten loci identified in (Rubin et al. 2022) and remaining after pruning for high LD between loci (see fig. S15). Below, the frequency of the largest allele in *G. fortis* in each of four species of ground-finches present on Daphne. In all cases, the largest species, *G. magnirostris,* also has the highest frequency of the large allele.

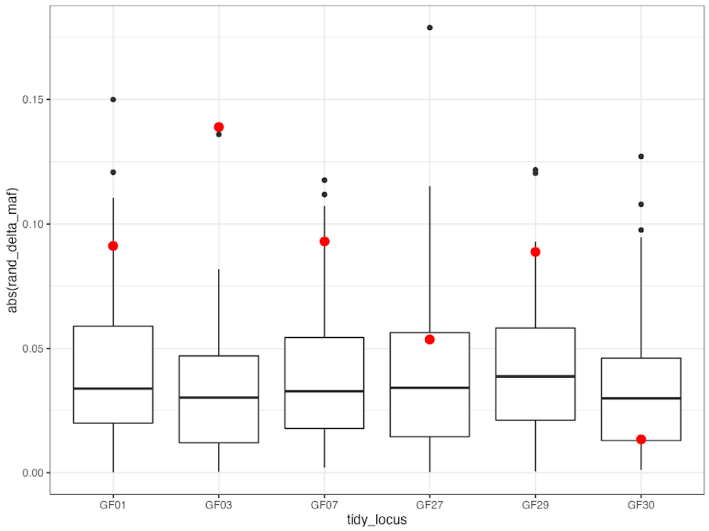


**fig. S21**: Boxplots show the change in allele frequency from 2004 to 2005 at 100 random SNPs with starting allele frequencies equal to those of the starting allele frequency for the focal locus. For four out of the six loci (in red), allele frequency changes at the identified six loci exceed those of randomly drawn loci.

**Table S1**: Summary of the number of samples used in the reference panel by species and island.

| **Genus** | **Species** | **Island** | **n** |
|---|---|---|---|
| *Camarhynchus* | *heliobates* | Isabela | 3 |
| *Camarhynchus* | *pallidus* | Fernandina | 3 |
| *Camarhynchus* | *pallidus* | Isabela | 4 |
| *Camarhynchus* | *pallidus* | San Cristóbal | 4 |
| *Camarhynchus* | *pallidus* | Santa Cruz | 5 |
| *Camarhynchus* | *parvulus* | Fernandina | 4 |

| | | | |
|---|---|---|---|
| *Camarhynchus* | *parvulus* | Floreana | 3 |
| *Camarhynchus* | *parvulus* | Isabela | 3 |
| *Camarhynchus* | *parvulus* | Santa Cruz | 12 |
| *Camarhynchus* | *parvulus* | Santa Fe | 3 |
| *Camarhynchus* | *pauper* | Floreana | 10 |
| *Camarhynchus* | *psittacula* | Fernandina | 5 |
| *Camarhynchus* | *psittacula* | Isabela | 4 |
| *Camarhynchus* | *psittacula* | Marchena | 2 |
| *Camarhynchus* | *psittacula* | Pinta | 9 |
| *Camarhynchus* | *psittacula* | Santa Cruz | 1 |
| *Certhidea* | *fusca* | Española | 10 |
| *Certhidea* | *fusca* | San Cristóbal | 13 |
| *Certhidea* | *olivacea* | Santiago | 5 |
| *Geospiza* | *acutirostris* | Genovesa | 4 |
| *Geospiza* | *bigbirds* | Daphne | 45 |
| *Geospiza* | *conirostris* | Española | 10 |
| *Geospiza* | *difficilis* | Fernandina | 4 |
| *Geospiza* | *difficilis* | Pinta | 10 |
| *Geospiza* | *difficilis* | Santiago | 4 |
| *Geospiza* | *fortis* | Albany | 3 |
| *Geospiza* | *fortis* | Daphne | 8 |
| *Geospiza* | *fortis* | Fernandina | 4 |
| *Geospiza* | *fortis* | Floreana | 4 |
| *Geospiza* | *fortis* | Isabela | 4 |
| *Geospiza* | *fortis* | Marchena | 5 |
| *Geospiza* | *fortis* | Pinta | 4 |

| | | | |
|---|---|---|---|
| *Geospiza* | *fortis* | Rabida | 2 |
| *Geospiza* | *fortis* | San Cristóbal | 6 |
| *Geospiza* | *fortis* | Santa Cruz | 10 |
| *Geospiza* | *fortis* | Santa Fe | 4 |
| *Geospiza* | *fortis* | Santiago | 2 |
| *Geospiza* | *fuliginosa* | Daphne | 4 |
| *Geospiza* | *fuliginosa* | Española | 4 |
| *Geospiza* | *fuliginosa* | Floreana | 4 |
| *Geospiza* | *fuliginosa* | Isabela | 4 |
| *Geospiza* | *fuliginosa* | Pinta | 4 |
| *Geospiza* | *fuliginosa* | San Cristóbal | 7 |
| *Geospiza* | *fuliginosa* | Santa Cruz | 4 |
| *Geospiza* | *fuliginosa* | Santiago | 10 |
| *Geospiza* | *hybrid* | Daphne | 15 |
| *Geospiza* | *magnirostris* | Daphne | 10 |
| *Geospiza* | *magnirostris* | Fernandina | 4 |
| *Geospiza* | *magnirostris* | Genovesa | 5 |
| *Geospiza* | *magnirostris* | Isabela | 4 |
| *Geospiza* | *magnirostris* | Marchena | 2 |
| *Geospiza* | *magnirostris* | Rabida | 4 |
| *Geospiza* | *magnirostris* | Santa Cruz | 4 |
| *Geospiza* | *propinqua* | Genovesa | 10 |
| *Geospiza* | *scandens* | Daphne | 4 |
| *Geospiza* | *scandens* | Floreana | 4 |
| *Geospiza* | *scandens* | Pinta | 4 |
| *Geospiza* | *scandens* | Rabida | 4 |

| | | | |
|---|---|---|---|
| *Geospiza* | *scandens* | San Cristóbal | 7 |
| *Geospiza* | *scandens* | Santa Cruz | 2 |
| *Geospiza* | *scandens* | Santiago | 4 |
| *Geospiza* | *septentrionalis* | Darwin | 4 |
| *Geospiza* | *septentrionalis* | Wolf | 8 |
| *Loxgilla* | *barbadensis* | Barbados | 5 |
| *Pinaroloxias* | *inornata* | Cocos | 8 |
| *Platyspiza* | *crassirostris* | Isabela | 4 |
| *Platyspiza* | *crassirostris* | Marchena | 4 |
| *Platyspiza* | *crassirostris* | Pinta | 4 |
| *Platyspiza* | *crassirostris* | San Cristóbal | 2 |
| *Platyspiza* | *crassirostris* | Santa Cruz | 5 |
| *Platyspiza* | *crassirostris* | Santiago | 4 |
| *Tiaris* | *bicolor* | Barbados | 3 |

**Table S2**: Post-hoc probabilities of pairwise differences between genotypes in mean beak length:

| Locus | Species | Major | AA v AB | AA v BB | AB v BB |
|---|---|---|---|---|---|
| G01 | *G. fortis* | BB | <0.0001 | <0.0001 | <0.0001 |
| G01 | *G. scandens* | AA | 0.0003 | 0.0036 | NS |
| G03 | *G. fortis* | BB | <0.0001 | <0.0001 | <0.0001 |
| G03 | *G. scandens* | BB | <0.0001 | <0.0001 | <0.0001 |
| G07 | *G. fortis* | BB | <0.0001 | <0.0001 | NS |
| G07 | *G. scandens* | AA | 0.0002 | NS | NS |

Major refers to the genotype with the largest mean beak length. NS = not significant ($P>0.05$).

### Supplementary Text

**Supplemental Text 1**: **Tests of the hypothesis of introgression from *G. fuliginosa* to *G. scandens* via *G. fortis*.**
Initial exploration of haplotype structure at the six loci revealed a third haplotype at locus *G03* that separate the two pointed haplotypes. We classified the pointed haplotype as either P1 (common in *G. scandens*) or P2 (common in *G. fuliginosa*) which enabled us to analyze variation at three haplotypes: B1, P1, and P2. *G. fortis*, *G. scandens* and *G. fuliginosa* differ markedly in haplotype frequencies. Differences are highly significant in the early part of the study prior to 1984 ($X^2$= 187.60, df 10, $P < 0.0001$) when the first (single) backcrossing was observed. Species differences at individual loci are greatest at P1P1 ($X^2$= 69.56), which is the commonest haplotype of *G. scandens* (0.76, $n$ = 33), at P2P2 ($X^2$= 68.07), which is the commonest haplotype of *G. fuliginosa* (0.65, $n$ = 43), and B1P2 ($X^2$= 14.73) and B1B1 ($X^2$= 24.14), which are the commonest haplotypes of *G. fortis* (0.36 and 0.32 respectively, $n$ = 85). All these $X^2$ values are significant ($P < 0.0001$).

The hypothesis is tested with three predictions. First, the prediction that *G. fuliginosa* is the source of introgressed genes into *G. fortis* is supported by a significant increase in the frequency of the P2P2 haplotype in *G. fortis* after 1984 ($F_{1,8}$ = 21.30, $P$ = 0.0017, adj $R^2$ = 0.69) (Fig. S1a). If *G. scandens* was an additional source, the frequency of P1P1 should increase also, but it did not change ($F_{1,8}$ = 2.97, $P$ = 0.1229, adj $R^2$ = 0.18). Second, the prediction that *G. fortis* is the source of introgressed genes into *G. scandens* is supported by the increase in frequencies of the combined haplotypes that contain the B-allele: B1B1, B1P1 and B1P2 ($F_{1,8}$ = 11.37, $P$ = 0.0098, adj $R^2$= 0.53) (S1b). Third, the prediction that *G. fuliginosa* is the source of genes in *G. scandens* received via the conduit of *G. fortis* is supported by an increase in the frequency of P2P2 in *G. scandens* ($F_{1,8}$ = 7.31, $P$ = 0.0269, adj $R^2$= 0.41; Fig. S1c).

**Supplemental Text 2**: **Origin of two *G. magnirostris* ancestries on Daphne**

These two ancestral populations reflect differences in morphology: one population is on average 2.4 mm larger (~5% of the smaller mean) in overall beak dimensions than the other (Welch's t-test, $t$ = 10.3, df = 196, $P < 0.0001$, Fig. S7). The smaller birds originated on one or more of the nearby islands of Santa Cruz, Marchena, and Isabela (testing six islands, n = 2-5 each, Table S1, Fig. S6). However, we could not determine an island of origin for the larger birds, which did not share ancestry with populations on any of the six islands we sampled. Of the remaining four unsampled populations (Santiago, Pinta, Wolf, and Darwin) the most likely candidates are those closest to Daphne, namely Santiago and Pinta. *G. magnirostris* are much larger on these two islands than on Wolf and Darwin (Grant et al. 1985).

**Supplemental Text 3: Genes contained in two QTLs**

*G01* contains a local peak overlapping *RNASEH2B,* which has been associated with chicken growth (Xie et al. 2012). The sixth locus, *G27*, is centered on the gene *LPP*. The sixth locus, *G27*, is centered on the gene *LPP*. Neither of these latter two loci intersect genes of known functional significance for morphology. This suggests that the nearest genes have an unknown role in craniofacial development or growth or the region contains long-range regulatory elements for physically distant genes.

**Supplemental Text 4**: **Comparisons to GCTA-LOCO**

We compared results generated in GEMMA to those generated in a leave-one-out-chromosome (LOCO) analysis in GCTA. The two analyses are both mixed models and allow us to include phenotypic covariates as well as sex and a measure of relatedness. Notably, the GCTA-LOCO analysis constructs the relatedness matrix by excluding the chromosome being analyzed. The analysis improves power when the QTL peaks are major drivers of the local relatedness matrix (Yang et al. 2014). We compared the results of the GCTA-LOCO analysis for the *G. fortis* cluster for beak PC1 and PC2, as well as body weight (Fig. S14). Although the same large effect loci were identified as in the GEMMA analysis, the regions of association tended to be substantially wider. In addition, chromosomes 3 and 5 suggest a large QTL (i.e. many megabases in length). The chromosome 5 region association in *G. fortis* is overlapping with the large region identified in the *G. scandens* GEMMA association analysis, which is a large region of high LD and low recombination. We interpret these broader association regions as indicative that excluding the chromosome being analyzed in the LOCO analysis inflates regions of broad LD, but recognize that additional QTLs may be present in these regions that the GEMMA analysis does not identify.

**Supplemental Text 5**: **Morphological associations at six loci in *G. fortis* and *G. scandens***

The six loci are associated with beak size in the same way in the three species (Fig. 3E). For beak shape the large allele at G01 and G07 is associated with a more pointed beak, while at G03 and G27 the large allele is associated with a blunter beak. To investigate the univariate basis of this difference we performed one-way ANOVAs separately on beak length and beak depth, which are the two major components of beak shape, and found that associations between mean beak length (but not beak depth) and genotype at G01 and G07 differ between *G. fortis* and *G. scandens*. Variation across genotypes at G01 and G07 runs in the opposite direction in *G. fortis* and *G. scandens* but in the same direction at other loci. Interspecific differences G07 are caused by the presence of multiple, ancestry-linked, haplotypes at these loci. At present it is not clear if this is due to multiple haplotypes with different phenotypic effects or epistatic interactions on different ancestry backgrounds at these loci.

*G. fortis* has a relatively blunt beak and *G. scandens* has a relatively elongated beak. In the main text, we focus on the PCA decomposition of three beak dimensions measured in the field. The purpose of this text is to examine how these differences in PCs are reflected in the three beak dimensions. We used one-way ANOVAs of beak length and beak depth, the major components of beak shape, to investigate how the two species vary in their associations between beak dimensions and genotypic variation at each of the six loci. We first excluded 307 hybrids and backcrosses determined from pedigrees, leaving a maximum of 1265 *G. fortis* and 610 *G. scandens* for analysis.

Variation in *G. fortis* exhibits a uniform pattern. At all six beak loci, LL genotypes have the largest mean beak length or depth, SS genotypes have the smallest and heterozygotes are intermediate. Each of the ANOVAs is significant at $P<0.0001$. *G. scandens* follows the same pattern in that the LL genotype is greater in beak depth than the SS genotype, albeit with less statistical support. For beak length, however, there are two noteworthy instances of a reversal of the size sequence. At G01 and G07, the sequence for mean beak length is AA > AB > BB in *G. scandens* ($P<0.0001$) whereas it is BB > AB > AA in *G. fortis* ($P<0.0001$). We tested pairwise differences at each of these loci by Tukey's HSD tests, and the results are given in Table S2. At each locus mean beak length differs significantly between the AA homozygote and the heterozygotes, but the species differ in the direction of the difference. This implies either heterogeneity at the loci and non-equivalence in the two species, or species-specific epistatic interactions with other loci.

The most likely explanation for these differences is epistatic interactions with other loci involved in beak length. The beak of *G. scandens* is considerably longer than the beak length in *G. fortis* on average and a large effect allele affecting beak depth need not necessarily reduce length. Evidence for this explanation can be found by considering the *G07* locus, where we discovered two pointed haplotypes: one is most common in *G. fuliginosa* and the other most common in *G. scandens* (Supplemental Text 1). The *G. fuliginosa* haplotype (P2) is associated with a shorter beak length than the *G. scandens* haplotype (P1), in both *G. fortis* (Tukey's $P < 0.0001$) and *G. scandens* (Tukey's $P < 0.0001$). The P2 haplotype is present at a greater frequency in *G. fortis* than *G. scandens* and when P2 is removed from the analysis, P1 is associated with the longer phenotype in both species. The P2 haplotype is also positively associated with genome-wide *G. fuliginosa* ancestry in both species (Supplemental Text 1), indicating that there is likely additional influence from other loci in the genome on beak length.

**References and Notes**


Alexander, D. H., J. Novembre, and K. Lange. 2009. Fast model-based estimation of ancestry in unrelated individuals. Genome Res. 19:1655–1664.

Aschard, H., B. J. Vilhjálmsson, A. D. Joshi, A. L. Price, and P. Kraft. 2015. Adjusting for heritable covariates can bias effect estimates in genome-wide association studies. Am. J. Hum. Genet. 96:329–339.

Danecek, P., J. K. Bonfield, J. Liddle, J. Marshall, V. Ohan, M. O. Pollard, A. Whitwham, T. Keane, S. A. McCarthy, R. M. Davies, and H. Li. 2021. Twelve years of SAMtools and BCFtools. Gigascience 10.

Delaneau, O., J.-F. Zagury, M. R. Robinson, J. L. Marchini, and E. T. Dermitzakis. 2019. Accurate, scalable and integrative haplotype estimation.

Enbody, E. D., C. G. Sprehn, A. Abzhanov, H. Bi, M. P. Dobreva, O. G. Osborne, C.-J. Rubin, P. R. Grant, B. R. Grant, and L. Andersson. 2021. A multispecies BCO2 beak color polymorphism in the Darwin's finch radiation. Curr. Biol. 31:5597–5604.e7.

Fox, J., and S. Weisberg. 2011. An {R} Companion to Applied Regression, Second Edition. SAGE publications, Thousand Oaks, CA.

Freed, D., R. Aldana, J. A. Weber, and J. S. Edwards. 2017. The Sentieon Genomics Tools - A fast and accurate solution to variant calling from next-generation sequence data.

Grant, P. R., and R. B. Grant. 2014. 40 Years of Evolution: Darwin's Finches on Daphne Major Island. Princeton University Press.

Kopelman, N. M., J. Mayzel, M. Jakobsson, N. A. Rosenberg, and I. Mayrose. 2015. Clumpak: a program for identifying clustering modes and packaging population structure inferences across K. Mol. Ecol. Resour. 15:1179–1191.

Korneliussen, T. S., A. Albrechtsen, and R. Nielsen. 2014. ANGSD: Analysis of next generation sequencing data. BMC Bioinformatics 15:356.

Korunes, K. L., and K. Samuk. 2021. pixy: Unbiased estimation of nucleotide diversity and divergence in the presence of missing data. Mol. Ecol. Resour. 1–10.

Lack, D. 1945. The Galapagos Finches (Geospizinae): A Study in Variation. California Acad. of Sciences.

Lamichhaney, S., J. Berglund, M. S. Almén, K. Maqbool, M. Grabherr, A. Martinez-Barrio, M. Promerová, C.-J. Rubin, C. Wang, N. Zamani, B. R. Grant, P. R. Grant, M. T. Webster, and L. Andersson. 2015. Evolution of Darwin's finches and their beaks revealed by genome sequencing. Nature 518:371–375.

Lamichhaney, S., F. Han, J. Berglund, C. Wang, M. S. Almén, M. T. Webster, B. R. Grant, P. R. Grant, and L. Andersson. 2016. A beak size locus in Darwin's finches facilitated character displacement during a drought. Science 352:470–474.

Lamichhaney, S., F. Han, M. T. Webster, L. Andersson, B. R. Grant, and P. R. Grant. 2018. Rapid hybrid speciation in Darwin's finches. Science 359:224–228.

Lawrence, M., W. Huber, H. Pagès, P. Aboyoun, M. Carlson, R. Gentleman, M. T. Morgan, and

V. J. Carey. 2013. Software for computing and annotating genomic ranges. PLoS Comput. Biol. 9:e1003118.
Li, H. 2013. Aligning sequence reads, clone sequences and assembly contigs with BWA-MEM. arXiv preprint arXiv 00:3.
Li, H., B. Handsaker, A. Wysoker, T. Fennell, J. Ruan, N. Homer, G. Marth, G. Abecasis, and R. Durbin. 2009. The Sequence Alignment/Map format and SAMtools. Bioinformatics 25:2078–2079.
Martin, M., M. Patterson, S. Garg, S. O. Fischer, N. Pisanti, G. W. Klau, A. Schöenhuth, and T. Marschall. 2016. WhatsHap: fast and accurate read-based phasing.
Meisner, J., and A. Albrechtsen. 2018. Inferring population structure and admixture proportions in low-depth NGS data. Genetics 210:719–731.
Picelli, S., A. K. Björklund, B. Reinius, S. Sagasser, G. Winberg, and R. Sandberg. 2014. Tn5 transposase and tagmentation procedures for massively scaled sequencing projects. Genome Res. 24:2033–2040.
Poplin, R., V. Ruano-Rubio, M. A. DePristo, T. J. Fennell, M. O. Carneiro, G. A. V. der Auwera, D. E. Kling, L. D. Gauthier, A. Levy-Moonshine, D. Roazen, K. Shakir, J. Thibault, S. Chandran, C. Whelan, M. Lek, S. Gabriel, M. J. Daly, B. Neale, D. G. MacArthur, and E. Banks. 2017. Scaling accurate genetic variant discovery to tens of thousands of samples. preprint at: https://www.biorxiv.org/content/10.1101/201178v3 201178.
Rastas, P. 2017. Lep-MAP3: robust linkage mapping even for low-coverage whole genome sequencing data. Bioinformatics 33:3726–3732.
Rezvoy, C., D. Charif, L. Guéguen, and G. A. B. Marais. 2007. MareyMap: an R-based tool with graphical interface for estimating recombination rates. Bioinformatics 23:2188–2189.
Rubinacci, S., D. M. Ribeiro, R. J. Hofmeister, and O. Delaneau. 2021. Efficient phasing and imputation of low-coverage sequencing data using large reference panels. Nat. Genet. 53:120–126.
Rubin, C.-J., E. D. Enbody, M. P. Dobreva, A. Abzhanov, B. W. Davis, S. Lamichhaney, M. Pettersson, C. Grace Sprehn, C. A. Valle, K. Vasco, O. Wallerman, B. Rosemary Grant, P. R. Grant, and L. Andersson. 2021. Darwin's finches - an adaptive radiation constructed from ancestral genetic modules.
Rubin, C.-J., E. D. Enbody, M. P. Dobreva, A. Abzhanov, B. W. Davis, S. Lamichhaney, M. Pettersson, A. T. Sendell-Price, C. G. Sprehn, C. A. Valle, K. Vasco, O. Wallerman, B. R. Grant, P. R. Grant, and L. Andersson. 2022. Rapid adaptive radiation of Darwin's finches depends on ancestral genetic modules. Science Advances 8:eabm5982.
Shringarpure, S. S., C. D. Bustamante, K. Lange, and D. H. Alexander. 2016. Efficient analysis of large datasets and sex bias with ADMIXTURE. BMC Bioinformatics 17:218.
Sprehn, C. G., E. Enbody, Y. Zan, and L. Andersson. 2021. Tn5 based tagmentation library prep protocol, high throughput v1. protocols.io.
Wood, S. N. 2011. Fast stable restricted maximum likelihood and marginal likelihood estimation of semiparametric generalized linear models. J. R. Stat. Soc. Series B Stat. Methodol. 73:3–36. Wiley.
Xie, L., C. Luo, C. Zhang, R. Zhang, J. Tang, Q. Nie, L. Ma, X. Hu, N. Li, Y. Da, and X. Zhang.

2012. Genome-wide association study identified a narrow chromosome 1 region associated with chicken growth traits. PLoS One 7:e30910.
Yang, J., A. Bakshi, Z. Zhu, G. Hemani, A. A. E. Vinkhuyzen, S. H. Lee, M. R. Robinson, J. R. B. Perry, I. M. Nolte, J. V. van Vliet-Ostaptchouk, H. Snieder, LifeLines Cohort Study, T. Esko, L. Milani, R. Mägi, A. Metspalu, A. Hamsten, P. K. E. Magnusson, N. L. Pedersen, E. Ingelsson, N. Soranzo, M. C. Keller, N. R. Wray, M. E. Goddard, and P. M. Visscher. 2015. Genetic variance estimation with imputed variants finds negligible missing heritability for human height and body mass index. Nat. Genet. 47:1114–1120.
Yang, J., S. H. Lee, M. E. Goddard, and P. M. Visscher. 2011. GCTA: a tool for genome-wide complex trait analysis. Am. J. Hum. Genet. 88:76–82.
Yang, J., N. A. Zaitlen, M. E. Goddard, P. M. Visscher, and A. L. Price. 2014. Advantages and pitfalls in the application of mixed-model association methods. Nat. Genet. 46:100–106.
Zhan, S., W. Zhang, K. Niitepõld, J. Hsu, J. F. Haeger, M. P. Zalucki, S. Altizer, J. C. de Roode, S. M. Reppert, and M. R. Kronforst. 2014. The genetics of monarch butterfly migration and warning colouration. Nature 514:317–321.
Zhou, X., and M. Stephens. 2014. Efficient multivariate linear mixed model algorithms for genome-wide association studies. Nat. Methods 11:407–409.
